# Exploring the Genetic Basis of Human Population Differences in DNA Methylation and their Causal Impact on Immune Gene Regulation

**DOI:** 10.1101/371872

**Authors:** Lucas T. Husquin, Maxime Rotival, Maud Fagny, Hélène Quach, Nora Zidane, Lisa M. McEwen, Julia L. MacIsaac, Michael S Kobor, Hugues Aschard, Etienne Patin, Lluis Quintana-Murci

## Abstract

DNA methylation is influenced by both environmental and genetic factors and is increasingly thought to affect variation in complex traits and diseases. Yet, the extent of ancestry-related differences in DNA methylation, its genetic determinants, and their respective causal impact on immune gene regulation remain elusive. We report extensive population differences in DNA methylation between individuals of African and European descent — detected in primary monocytes that were used as a model of a major innate immunity cell type. Most of these differences (~70%) were driven by DNA sequence variants nearby CpG sites (meQTLs), which account for ~60% of the variance in DNA methylation. We also identify several master regulators of DNA methylation variation in *trans*, including a regulatory hub nearby the transcription factor-encoding *CTCF* gene, which contributes markedly to ancestry-related differences in DNA methylation. Furthermore, we establish that variation in DNA methylation is associated with varying gene expression levels following mostly, but not exclusively, a canonical model of negative associations, particularly in enhancer regions. Specifically, we find that DNA methylation highly correlates with transcriptional activity of 811 and 230 genes, at the basal state and upon immune stimulation, respectively. Finally, using a Bayesian approach, we estimate causal mediation effects of DNA methylation on gene expression in ~20% of the studied cases, indicating that DNA methylation can play an active role in immune gene regulation. Using a system-level approach, our study reveals substantial ancestry-related differences in DNA methylation and provides evidence for their causal impact on immune gene regulation.

## Introduction

Individuals and populations display variable susceptibility to infectious diseases, chronic inflammatory disorders, and autoimmunity [1, 2]. Over the last decade, it has become clear that such disparities partly result from differences in the host genetic make-up, with an increasing number of genes accounting for varying abilities to fight infections at the individual and population level [3, 4]. Furthermore, population genetic studies have revealed that pathogen-driven selection has substantially impacted human genetic diversity [5, 6]. Because the mortality, and thus the selective pressure, imposed by pathogens have been paramount [7], human populations had to adapt to the different pathogenic environments they encountered around the globe, and genes involved in host defence are among the functions most strongly targeted by natural selection [5, 8-11]. While substantial evidence supports this hypothesis at the genetic level, we still know little about the degree of naturally-occurring epigenetic variation at the population level and how this may impact immune phenotypes.

As the immune system is the primary interface with the human pathogenic environment, the study of DNA methylation [12, 13] offers a unique opportunity to explore the interplay between the genome and environmental cues. DNA methylation can be affected by a range of external factors, such as nutrition, toxic pollutants, social environment and infectious agents [14-19]. Furthermore, numerous studies have mapped DNA sequence variants associated with DNA methylation variation [20-28], i.e., methylation quantitative trait loci (meQTLs), and ~20% of the inter-individual variation in DNA methylation has been attributed to genetic factors [29, 30]. DNA methylation variation has also been associated with complex traits, including aging [31], body mass index [32], various cancers [33, 34], obesity [35], as well as autoimmune and inflammatory disorders [36, 37]. Yet, most studies of human epigenome variation, both in health and disease conditions, have focused on populations of homogeneous genetic ancestry, primarily of European-descent.

A few studies, however, have reported that population differences in ancestry, habitat or lifestyle affect DNA methylation, providing an initial assessment of the contribution of genetic factors and gene-environment (G×E) interactions to population-level epigenetic variation [38-44]. Yet, these studies investigated DNA methylation variation from virus-transformed lymphoblastoid cell lines or whole blood, so the differences observed could reflect, at least partially, epigenetic changes induced by cell immortalization or heterogeneity in blood cell composition that was not fully accounted for [45-47]. Thus, the extent of DNA methylation variation related to ancestry, and its genetic determinants, in a cellular setting relevant to immunity are far from clear.

A growing body of research has reported ancestry-related variation in terms of immune gene expression levels. Two recent studies found marked differences between individuals of African and European ancestry in their transcriptional responses to infectious challenges [48, 49], and showed that regulatory variants (i.e., expression quantitative trait loci, eQTLs) explain a substantial proportion of these population differences. Still, a substantial fraction of the variance in gene expression, both across individuals and populations, cannot be attributed to genetic factors and remains unexplained [48-55]. In this context, DNA methylation represents an additional, possible layer for variation in gene regulation [56]. The observed correlations between DNA methylation and gene expression levels can be positive and negative; in the canonical model, high levels of methylation at promoter regions is often associated with low gene expression, but elevated gene body methylation is also associated with active expression [28, 47, 57-60]. There is also increasing evidence that DNA methylation can play both passive and active roles in the regulatory interactions influencing gene expression, but the causality relationships between DNA methylation, gene expression and genetic factors are not fully understood [19, 23, 56]. Furthermore, genetic variants associated with complex traits or diseases by genome-wide association studies (GWAS) often overlap both eQTLs and meQTLs, suggesting that disease risk can be mediated, directly or indirectly, by variation in DNA methylation [61-67].

Here, we aimed to broad our understanding of the mechanistic links between ancestry-related differences in DNA methylation, genetic factors and immune gene regulation. To do so, we build upon the EvoImmunoPop collection of primary monocytes originating from 200 healthy western Europeans of self-reported African and European ancestry [48]. We generated high-density DNA methylation profiles using the Infinium MethylationEPIC array, which captures methylation variation at more than 850,000 sites. This new dataset was combined with both genome-wide genotyping and whole-exome sequencing data, as well as with RNA-sequencing profiles from resting and stimulated monocytes with various immune stimuli, obtained from the same individuals. Such a system-level approach, integrating epigenetic, genetic and transcriptional data, allowed us to assess the extent to which population-level variation in DNA methylation and its genetic determinants impact transcriptional activity related to immune responses.

## Results

### Population differences in DNA methylation profiles of primary monocytes

To assess population differences in DNA methylation of a purified innate immunity cell type, we characterized DNA methylation variation at > 850,000 CpG sites across the genome, in monocytes originating from individuals of African descent (AFB) and European descent (EUB) all living in Belgium. After normalization and filtering (see “**Methods**”), we retained a final dataset of 552,141 methylation sites in 156 individuals (78 of each ethnic group, **Figure S1**). Principal component analysis (PCA) of DNA methylation clearly separated AFB and EUB along the first two PCs, which explained together 11.6% of the total variance (**Fig. 1a**). At a false discovery rate (FDR)=1%, we identified 77,857 sites (14.1% of the total number) that presented a significant difference between AFB and EUB in their mean level of DNA methylation. When restricting our analyses to CpGs that presented a mean difference > 5% (measured by the *β*-value [68], see “**Methods**”), we identified a total of 12,050 differentially methylated sites between populations (DMS) that mapped to 4,818 genes.

**Fig. 1.**
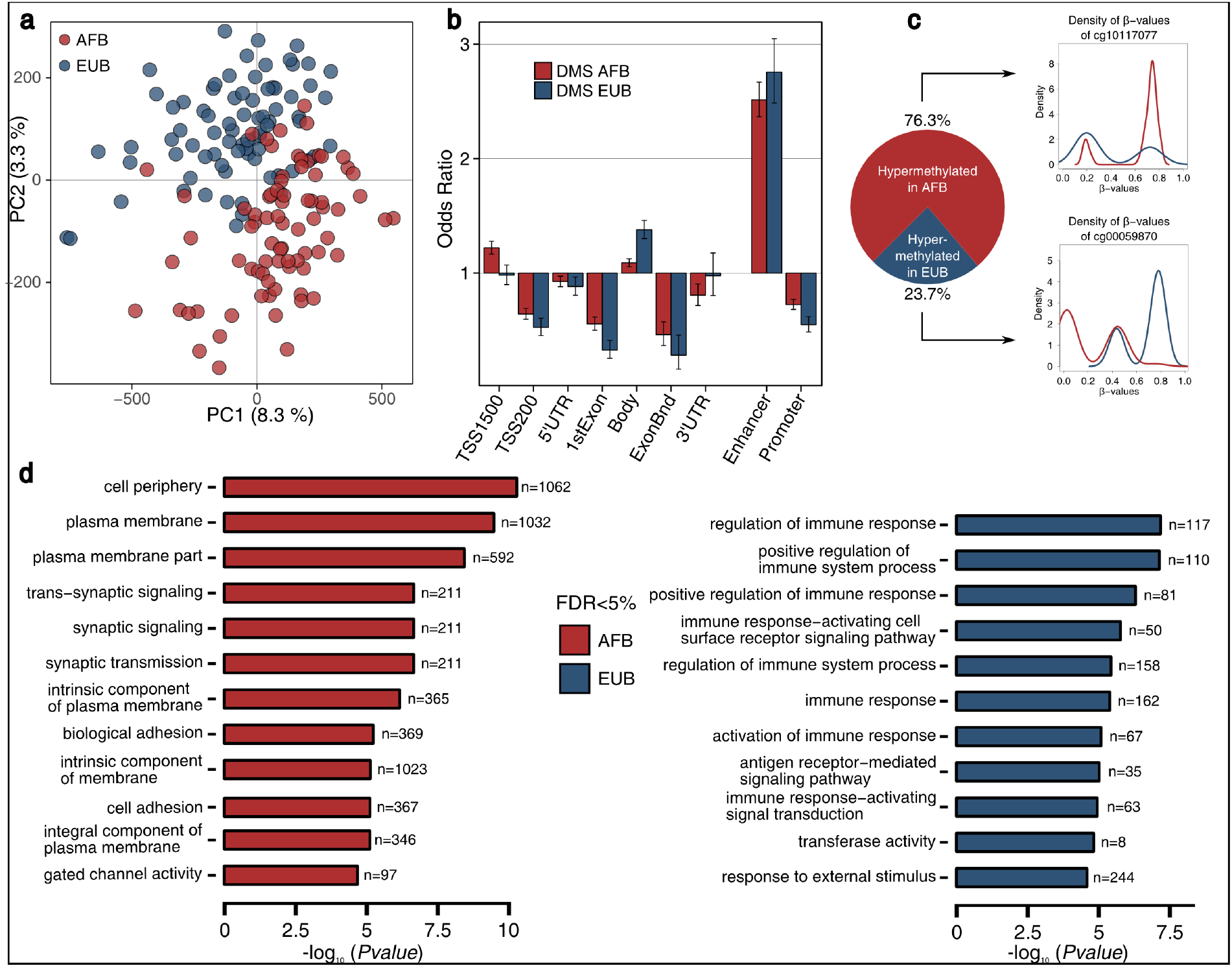
Population differences in DNA methylation profiles. **a** Principal Component Analysis (PCA) of DNA methylation profiles for all 156 individuals. Red and blue circles represent African (AFB) and European (EUB) individuals, respectively. The proportions of variance explained by PC1 and PC2 are indicated. **b** Genomic location of differentially methylated sites (DMS), for CpG sites hyper-methylated in AFB (red) and in EUB (blue). Odds ratio and 95% confidence intervals are displayed for AFB-DMS and EUB-DMS, comparing their localization in different genomic locations as provided by Illumina (TSS1500, TSS200, 5’UTR, 1stExon, Body, Exon boundaries [ExonBnd] and 3’UTR), and in enhancer and promoter regions specifically detected in monocytes by ChromHMM phase 15 (see refs. [106, 107]). Odds ratios were computed against the general distribution of the 552,141 CpGs of our dataset. **c** Proportion of DMS that are either hypermethylated in AFB (red) or in EUB (blue) individuals. The density of β-values of one CpG site by category is given as an illustration of the population differences, with red and blue lines representing the methylation density in AFB and EUB, respectively. **d** Gene Ontology (GO) enrichment analyses of AFB- and EUB-DMS. For both groups, the top-GO categories reaching 5% FDR are shown, together with the number of genes per category and the log-transformed FDR-adjusted enrichment *P*-values.

The genomic distribution of DMS, which were highly enriched in enhancer regions (Odds ratio (OR) ~2.6, *P* = 1.42×10^−224^), was independent of the population where hyper-methylation was observed (**Fig. 1b**). However, of the 12,050 DMS, 76.3% were more methylated in AFB than in EUB, with respect to the observed 54% when considering all CpGs (Fisher’s exact *P* < 2.2×10^−16^) (**Fig. 1c)**, and the corresponding genes were enriched in Gene Ontology (GO) categories related to cellular periphery and plasma membrane (**Fig. 1d**). The remaining 23.7%, which were hyper-methylated in EUB, were enriched in sites located in genes largely associated with immune response regulation and responses to external stimulus (**Fig. 1c, d**; **Table S1**). These results, which cannot be explained by population differences in monocyte subpopulations (i.e. CD14_high_/CD16_neg_ [Classical], CD14_high_/CD16_low_ [Intermediate] and CD14_low_/CD16_high_ [Non-Classical]), reveal genes and functions that present extensive population differences in DNA methylation in primary monocytes.

### Genetic factors drive most ancestry-related DNA methylation variation

We next examined the genetic determinants of the DNA methylation differences observed, and mapped methylation quantitative trait loci (meQTLs). We first tested for local associations between DNA methylation variation at CpGs and SNPs located within a 100-kb window (*cis*-meQTLs), using MatrixEQTL [69] (see “**Methods**”). We set a 5% FDR threshold, considering one association per CpG site and using 100 permutations (*P <* 1×10^−5^). We adjusted for age, surrogate variables (i.e. known batch effects and unknown confounders), and the first two PCs of the genetic data (**Figure S2**), to account for population stratification. To detect subtle effects, we merged all individuals and included ancestry as a covariate, but, simultaneously, we analysed the two populations separately to detect population-specific effects. For all subsequent analyses, we present the significant results of these two approaches combined, unless otherwise indicated.

We identified 69,702 CpGs associated with at least one genetic variant in at least one population (~12.6% of all sites, referred to as meQTL-CpGs). Given that multiple linked SNPs can be associated to the same CpG, we kept the best-associated SNP for each meQTL-CpG. However, we also used a fine mapping approach [51] to detect independent SNPs associated to each CpG (see “**Methods**”). In doing so, we detected 9,826 additional meQTLs (**Figure S3**), providing a more thorough view of the contribution of proximate genetic variants to DNA methylation variation. The median distance between a CpG and its associated SNP was ~3.8 kb (**Figure S4**), supporting the close genetic control of DNA methylation [22, 28, 41, 65]. Furthermore, we found a 2.2-fold enrichment of meQTL-CpGs in enhancers (*P* < 1×10^−326^), a trend that was even more pronounced for meQTLs associated with population differences in DNA methylation (meQTL-DMS; OR ~2.8, *P* = 6.8×10^−317^, **Figure S5**).

Focusing on ancestry-related differences, we observed that ~70.2% of DMS harbour a significant meQTL, with respect to the 12% detected genome-wide (Fisher’s exact *P* < 2.2×10^−16^; **Fig. 2a**). These meQTLs were found to account, on average, for ~58% of the observed population differences in DNA methylation (**Figure S6**, see “**Methods**”). Furthermore, they presented opposite effects on DNA methylation as a function of their population differences in allelic frequency; i.e. a derived allele at higher frequency in Africans was associated with high levels of DNA methylation, while the opposite was observed for meQTLs at higher frequency in Europeans (**Fig. 2b**). This observation provides a genetic explanation for the unbalanced patterns of hyper-methylation, observed at DMS, between Africans and Europeans (**Fig. 1c**)

**Fig. 2.**
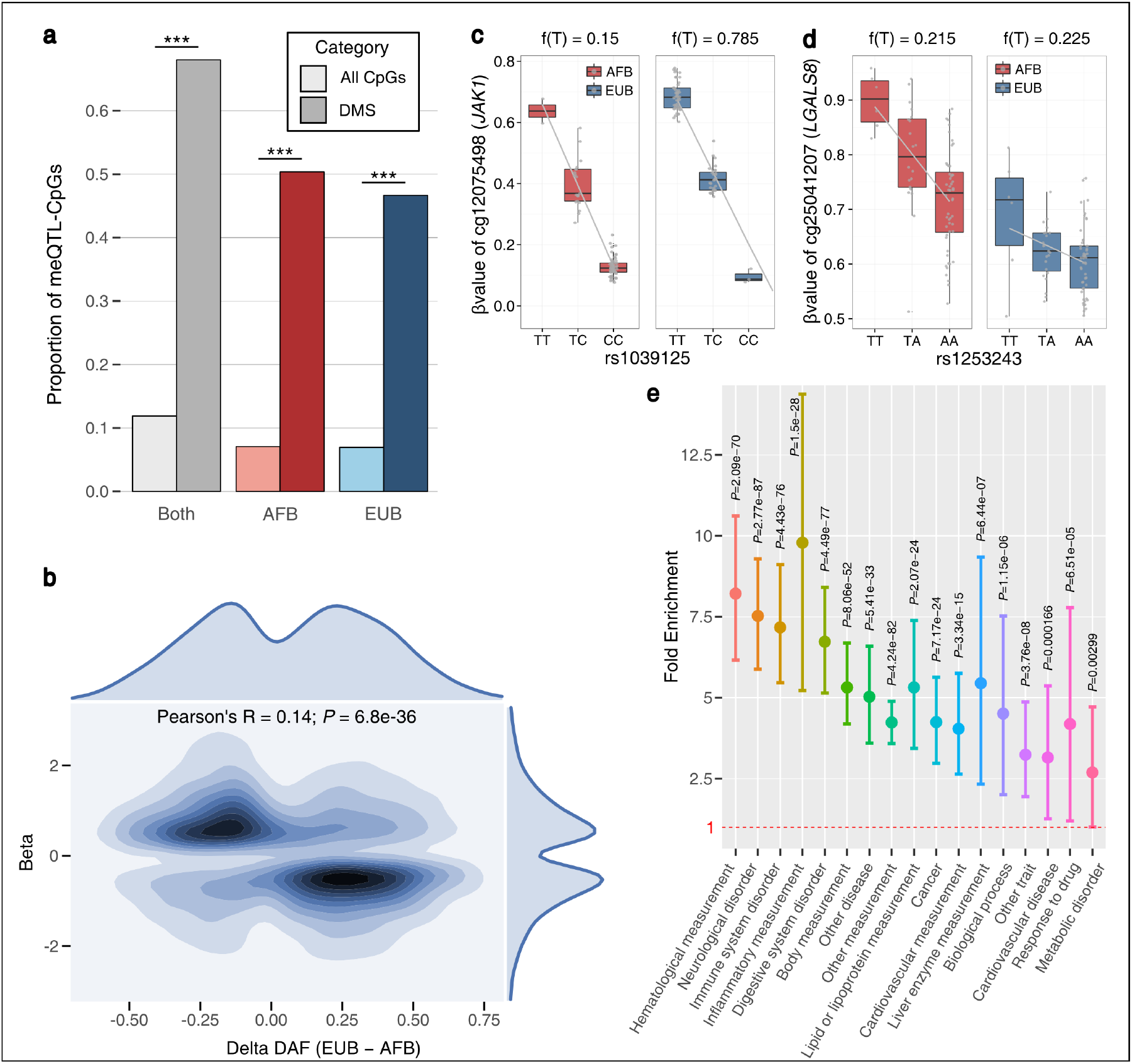
Genetic control of population differences in DNA methylation levels. **a** Proportions of CpGs and DMS associated to genetic variants identified in the three meQTL studies: merging the two populations (grey shades), mapping in AFB only (red shades) and in EUB only (blue shades). For each mapping, proportions among all 552,141 tested CpG sites, and among DMS, are indicated in light and dark colours, respectively. *** Fisher’s exact *P* < 2.2×10^−16^. **b** Contour plot of meQTL effects on DMS as a function of their difference in derived allelic frequencies (DAF) between populations. For each of the 8,459 DMS for which we detected at least one meQTL, we used a Kernel Density Estimation to draw the contour plot of the effect of the derived allele of the meQTL onto methylation (Beta, Y axis) according to the ∆DAF (DAF_EUB_ – DAF_AFB_, X axis). The coefficient and *P*-value of the Pearson’s correlation test are displayed. The marginal distribution of the two variables is displayed: top for ∆DAF, and right for Beta. **c-d** Examples of meQTLs detected in this study. Boxplots represent the distribution of β-values as a function of genotype, for AFB (red) and EUB (blue) individuals. The minor allele frequency of each meQTL is presented for each population on the top. Grey lines indicate the fitted linear regression model for β-value~genotype for each population. **e** Fold enrichment of meQTLs associated with DMS in GWAS hits. For each of the 17 parental EFO categories, the fold enrichment, the 95% confidence intervals obtained by bootstrap and the associated *P* values are shown.

Local meQTLs can, a priori, lead to population differences in DNA methylation following two main models: (i) the meQTL has a similar effect in both populations but present different allelic frequencies (**Fig. 2c**), or (ii) the meQTL is present at similar frequencies but display population-specific effects, revealing more complex interactions (**Fig. 2d**). While 18,250 and 17,572 meQTL-CpGs were detected exclusively in AFB and EUB, respectively, 33,880 were detected in both populations. Among the latter, we sought to identify population differences in the intensity of the association, by fitting, for each meQTL-CpG, a linear model including an interaction term between population and each independent genetic effect. In doing so, we detected 1,467 significant population-specific effects, supporting the occurrence of G×E or G×G effects.

### Ancestry-related meQTLs are enriched in associations with complex traits and diseases

Given that a large fraction of genetic variants identified by GWAS are thought to act by affecting gene regulation [70-73], we investigated the putative functional impact of the detected meQTLs on ultimate complex phenotypes. In practice, we searched for enrichments in GWAS hits among our set of 79,528 meQTLs, correcting for linkage disequilibrium (see “**Methods**”). Focusing on the 17 parental classes of the Experimental Factor Ontology (EFO) classification [74], we found that meQTLs were enriched in significant hits for all these functional categories (**Figure S7**, OR ~2.1-5.5, *P* < 4.1×10^−10^). Stronger enrichments were detected for meQTLs associated with population differences in DNA methylation (OR ~2.7-9.8, *P* < 2.9×10^−3^), in particular for phenotypes related to haematological measurements, neurological disorders, immune system disorders, inflammatory measurements and digestive system disorders (**Fig. 2e**).

Because DNA methylation and meQTLs have been shown to be largely cell or tissue dependent [23, 75-80], we next searched for the specific traits that account for the signals detected at the parental category “immune system disorder”, given our focus on primary monocytes. We found that meQTLs overlapped variants associated with diseases such as osteoarthritis, psoriasis, systemic lupus erythematosus, inflammatory skin disease or type 1-diabetes (**Figure S8**). For example, the meQTL SNP rs629953 presents markedly different frequencies between AFB and EUB (DAF AFB 7.5% *versus* DAF EUB 62%), leading to variable population-level DNA methylation at *TNFAIP3* (cg06987098), and has been previously associated with psoriasis susceptibility [81, 82]. Together, our analyses support that complex traits and variable DNA methylation are pleiotropically associated with genetic variation [39, 60, 63, 64], but extend these associations to variants affecting ancestry-related epigenetic variation in the context of an innate immunity cell type.

### Exploring the distant genetic control of DNA methylation variation

We subsequently searched for the effects of distant genetic variants on DNA methylation variation (*trans*-meQTLs). To limit the burden of multiple testing, we focused on 73,561 SNPs located nearby (+/-10kb of the TSS) 600 genes encoding transcription factors (TF), because *trans*-meQTLs are enriched in *cis*-eQTLs for TF-coding genes [65]. Only associations for which the SNP-CpG distance was higher than 1 Mb were considered, at an FDR of 5% (*P <* 1×10^−9^). Given the generally low power to map *trans*-associations, we performed this analysis by considering all individuals together and including ancestry as a covariate.

We identified 102 CpG sites associated with at least one distant SNP, for a total of 483 *trans*-meQTLs that involved 79 independent loci (**Table S2**). Among these, we detected a number of hubs of distant genetic control of DNA methylation variation, including five TFs (*CTCF*, *FOXI1*, *ZBTB25*, *MKL2* and *NFATC1*) where local genetic variation was associated with at least 10 different CpGs in *trans* (**Table S2**). Highlighting one pertinent example, a single genetic variant (rs7203742) nearby *CTCF* — encoding a transcriptional regulator with 11 highly conserved zinc-finger domains — controls the degree of DNA methylation at 30 CpG sites, ~29.4% of all CpGs regulated in *trans*. Furthermore, of the 21 *trans*-regulated CpGs that were detected as DMS, 12 were controlled by the same *CTCF* variant. That this variant (T*→*C) presents high levels of population differentiation (DAF AFB 24% *vs*. EUB 88%, *F*_ST_=0.59 in the 1% of the genome-wide distribution) suggests the action of positive selection targeting the derived allele in Europeans. This observation makes of *CTCF* not only a master regulator of DNA methylation, as previously observed [65], but also an important contributor to differences in DNA methylation between human populations.

### Dissecting the mechanistic relationships between DNA methylation and gene expression

We leveraged the availability of RNA-sequencing data from the same individuals [48] to obtain new insights into the mechanistic relationships between DNA methylation and gene expression variation, in African and European individuals. We associated the levels of expression of 12,578 genes in primary monocytes with those of DNA methylation at CpGs located within 100 kb of their TSS, for a total of 513,536 CpG sites. Associations were considered significant if they passed a *P*-value threshold determined using 100 permutations (FDR=5%, *P* < 5×10^−5^) (see “**Methods**”).

We identified 1,666 CpGs whose levels of DNA methylation were associated with gene expression (eQTMs), for a total of 811 genes (eQTM-genes) associated with at least one CpG in one population group (**Table S3**). The KEGG pathways associated with eQTM-genes contained a large number of immune-related pathways, providing a link between DNA methylation and gene expression in the context of immunity (**Fig. 3a**). While we detected 136 and 168 eQTM-genes specifically in AFB and EUB, partially reflecting population differences in gene expression and DNA methylation variance, the vast majority (62%) were shared between populations. To identify ancestry-related effects among the 507 shared eQTM-genes, we fitted a linear model including an interaction term between population and each independent epigenetic effect, and found 25 eQTM-genes where the intensity of the association differed between populations. That these 25 cases all corresponded to genes whose eQTMs were also under genetic control suggests, again, the occurrence of G×G or G×E interactions.

**Fig. 3.**
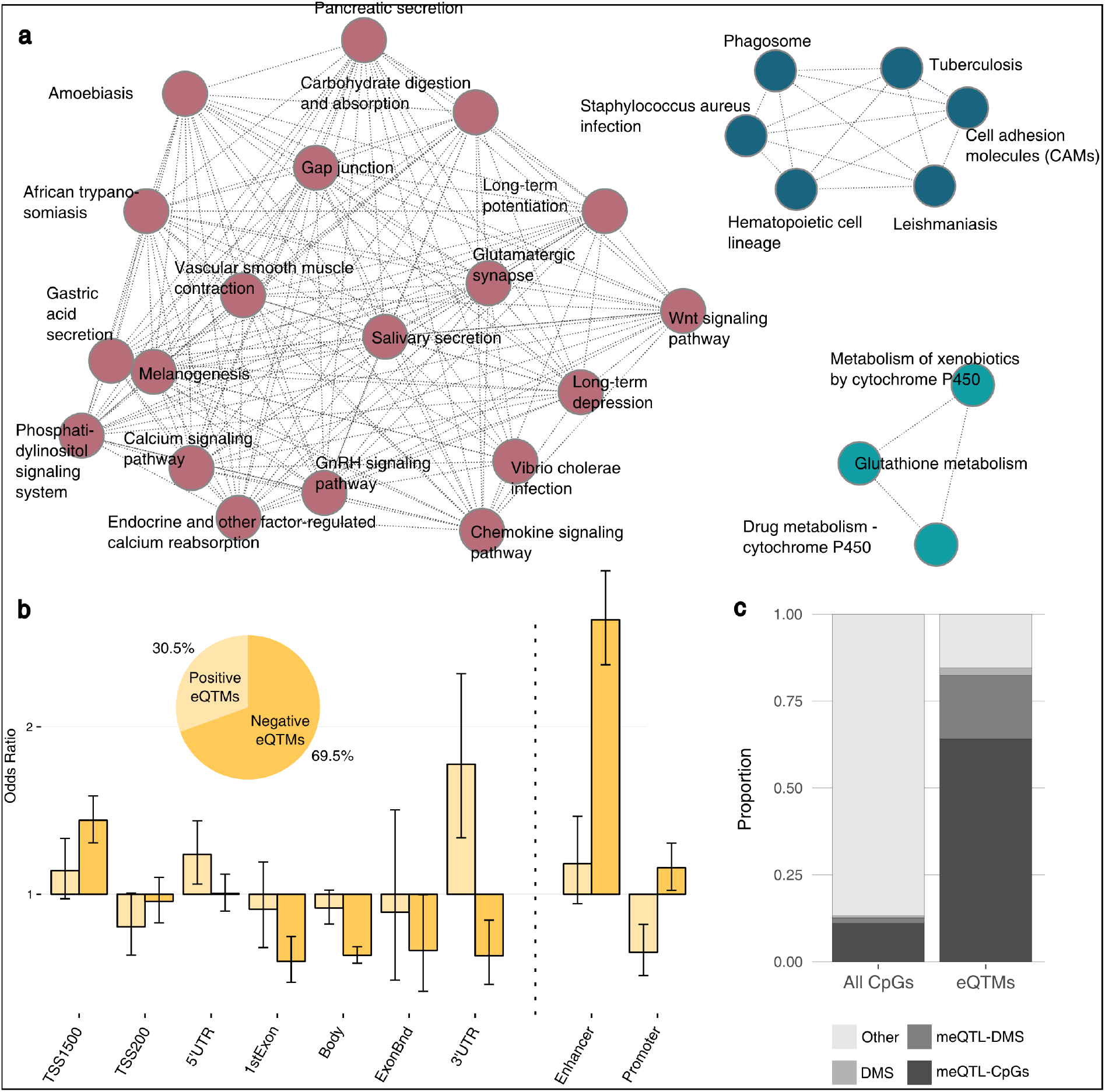
Correlations of DNA methylation with gene expression. **a** Networks of KEGG pathways of genes detected in the eQTM mapping. **b** Genomic location of eQTMs, for positively and negatively associated CpG sites (light and dark yellow, respectively). Odds ratio were computed against the general distribution of the 552,141 CpGs from our dataset. The distribution of eQTMs according to the direction of their effect on gene expression is shown. **c** Proportions of different groups of CpG sites in all tested sites (left panel) and among the detected eQTMs (right panel).

Based on current genomic annotations, eQTMs were mostly negatively correlated to gene expression (69.5% *vs.* 30.5%, see also refs. [23, 28, 65, 83, 84]). Negatively correlated sites were strongly enriched in enhancers (OR~2.6, *P =* 6.6×10^−59^) (**Fig. 3b**), highlighting their major role in transcriptional regulation [85-87]. In addition, we found a slight excess of negative associations in promoters (OR ~1.2, *P =* 1.8×10^−2^) and nearby TSS (TSS1500) (OR ~1.4, *P =* 7.2×10^−13^), as expected following the canonical model. Conversely, positive associations were enriched in sites located nearby UTRs, particularly 3’-UTR (OR ~1.8, *P =* 8.4×10^−5^) [88], but depleted in sites located in promoters (OR ~0.6, *P =* 1.1×10^−4^) (**Fig. 3b**). Furthermore, we found that eQTMs were strongly enriched in DMS (OR ~11.8, *P* < 1.93×10^−216^) and, importantly, in meQTL-CpGs (OR ~33.2, *P* < 1×10^−326^) (**Fig. 3c**). Together, these observations indicate that DNA methylation variation, in particular at sites that are differentially methylated across populations (DMS), are much more likely to be under genetic control when associated with gene expression differences (eQTMs), than random CpG sites.

### Exploring the underlying causality between regulatory loci and gene expression

Because the respective roles of genetic and epigenetic factors in transcriptional regulation are not fully understood [56], we next mapped eQTLs (FDR=5%, see “**Methods**”) to identify the situations where DNA methylation, gene expression and genetic variants show significant associations between all pairs (**Figure S9**). We thus obtained 552 trios, each of them consisting of one gene, one to various CpGs and one to various SNPs (containing 68.1% of the genes detected in the eQTM mapping). This suggested potential, causal relationships between these variables — a latent, though challenging, question in epigenetics. To infer causality between regulatory loci (i.e. eQTMs and eQTLs) and gene expression variation for these specific trios, we first used an elastic net model to build two intermediate variables measuring (i) DNA methylation variability attributable to genetics for the trios presenting more than one SNP, and (ii) gene expression variability attributable to DNA methylation for the trios presenting more than one CpG (see “**Methods**”).

We used a Bayesian approach [89] to assess potential causal effects of a mediating variable *M* (DNA methylation) on the relationship between an independent variable *X* (genetics) and a dependent variable *Y* (gene expression) [90]. When comparing the performance of this method with that of an approach based on partial correlations, using simulated data and various genomic scenarios, we found similar results between the two approaches in terms of sensitivity and specificity (**Fig. 4a-b; Figure S10;** see **“Methods”**). We then ran the mediation analysis on each trio, adjusting for regular covariates (age and surrogate variables), but also for the 4^th^ and 2^nd^ PCs of gene expression and DNA methylation, respectively. The latter covariates were added because they likely capture potential confounding factors inducing correlation between DNA methylation and expression, which would violate the assumption of the causal inference model (**Figure S11**). Note that reverse causation was found to be unlikely in our experimental setting and was thus not considered in our analyses (**Supplementary Note 1**).

**Fig. 4.**
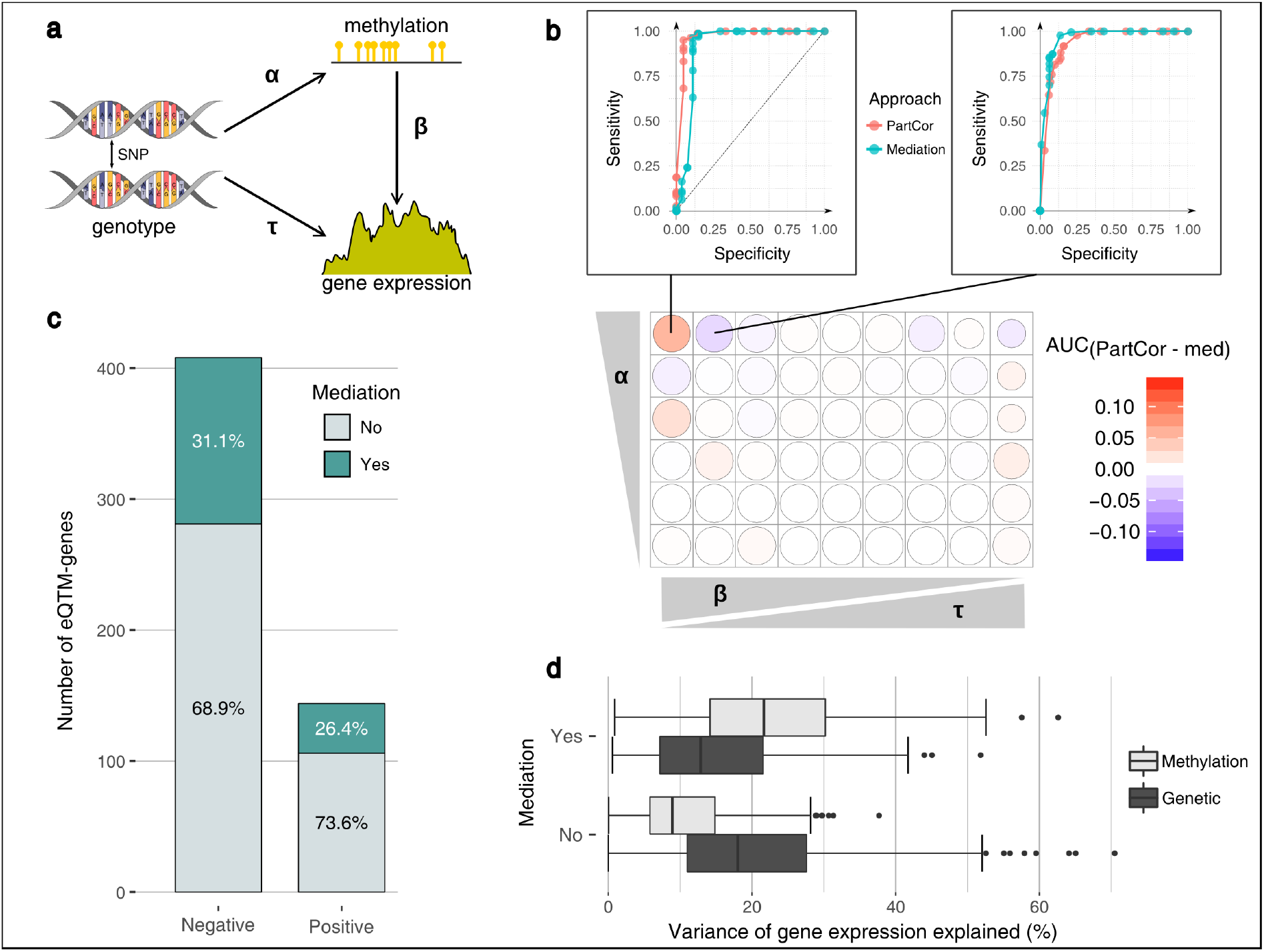
Inference of the causal effects of DNA methylation on gene regulation. **a** Representation of a simulated scenario, with the three varying parameters (α, β and τ). **b** Comparison of the mediation analysis (med) with a partial correlation approach (PartCor) using a range of different simulated parameters for α (0.3-0.8), β (0.9-0.1) and τ (0.1-0.9). Note that the parameter range simulated for β and τ was adjusted so that we kept 75% of the variance unexplained (random noise parameter γ=0.25). The difference of the area under the curve (AUC) between the two approaches is represented with different shades of red and blue. The sizes of the circles are proportional to the mean AUC of the two approaches. Two examples of the ROC curves are shown in the upper part of the figure. **c** Number of mediated and non-mediated eQTM-genes for negative and positive associations between DNA methylation and gene expression. The percentages of these two categories are also indicated. **d** Proportion of variance of gene expression explained by DNA methylation (light gray) and genetics (dark gray), in mediated and non-mediated cases.

At FDR=5%, we identified 165 genes where the genetic control of expression levels was mediated by DNA methylation (i.e., α×β was significantly different from zero, **Fig. 4a**), in at least one population. Remarkably, in 66 of these cases, mediation occurred through CpG sites that are differentially methylated across populations (DMS) (**Table S4**). The proportion of mediated genes whose expression was positively and negatively correlated to DNA methylation was similar, ranging from 26% to 31% (**Fig. 4c**). Expectedly, we found that, among mediated genes, DNA methylation explained a significantly higher proportion of the variance of gene expression than genetics (mean R^2^= 23.4% versus 15.4%, respectively; Wilcoxon *P* = 3.3×10^−11^), in contrast with the 387 non-mediated cases where we observed the opposite trend (Wilcoxon *P* = 7.8×10^−37^) (**Fig. 4d**).

We also found that CpG sites mediating gene expression were preferentially located in enhancers (OR ~2.5, *P* = 4.0×10^−21^), highlighting again the major role of these regions in epigenetic regulatory mechanisms [91-93]. These CpGs were depleted in promoters (OR ~0.7, *P* = 1.4×10^−2^), which were otherwise enriched in non-mediating CpGs (OR ~1.3, *P* = 5.9×10^−3^). Among mediated cases, we found key genes of the immune response, such as *NLRP2*, *RAI14*, *NCF4* or *ICAM4*, and, interestingly, genes with functions related to transcriptional activity, encoding zinc-finger proteins (**Table S4**). This suggests a more extensive role of DNA methylation in regulating gene expression than the local associations described here, through the regulation of DNA-binding protein activity.

### Impact of immune perturbation on genetic and epigenetic interactions

Finally, we sought to understand how DNA methylation variation at the basal state affects transcriptional responses to immune activation. We used RNA-sequencing data, obtained from the same individuals, after exposure to various stimuli: LPS activating TLR4 and Pam3CSK4 activating TLR1/2, both pathways sensing bacterial components, R848 activating TLR7/8, predominantly sensing viral nucleic acids, and influenza A virus (IAV) [48]. We then mapped response-QTMs (reQTMs) using fold-changes in gene expression between non-stimulated and stimulated states, for all genes expressed in either conditions (see “**Methods**”).

We found 230 genes whose response to immune activation was associated with DNA methylation in at least one condition; most associations were context-specific, with only 7 genes detected in all conditions (**Fig. 5a; Table S3**). Furthermore, a 2.5-fold increase was observed in the number of reQTM-genes detected upon activation with viral-stimuli (R848 and IAV; 197 unique genes) with respect to those detected for bacterial ligands (LPS and Pam3CSK4; 78 unique genes) (**Fig. 5a**). For example, we detected a reQTM upon R848 stimulation for *CARD9* in EUB and *CD1D* upon IAV infection in AFB, both genes known to play an important role in host defence (**Fig. 5b-c**). Despite reQTMs and eQTMs present a similar genomic distribution (**Figure S12**), we observed an important shift towards positive associations between DNA methylation and transcriptional responses, in particular to TLR ligands (**Fig. 5d**). However, two distinct groups of reQTMs were apparent: reQTMs that present the strongest associations between DNA methylation and gene expression at the stimulated state (**Fig. 5b**), and reQTMs that present the strongest associations in the non-stimulated condition (**Fig. 5c**). We found that the general shift towards positive associations was mainly accounted by the latter group, with associations between DNA methylation and expression upon stimulation following primarily the canonical model of negative associations (**Figure S13**).

**Fig. 5.**
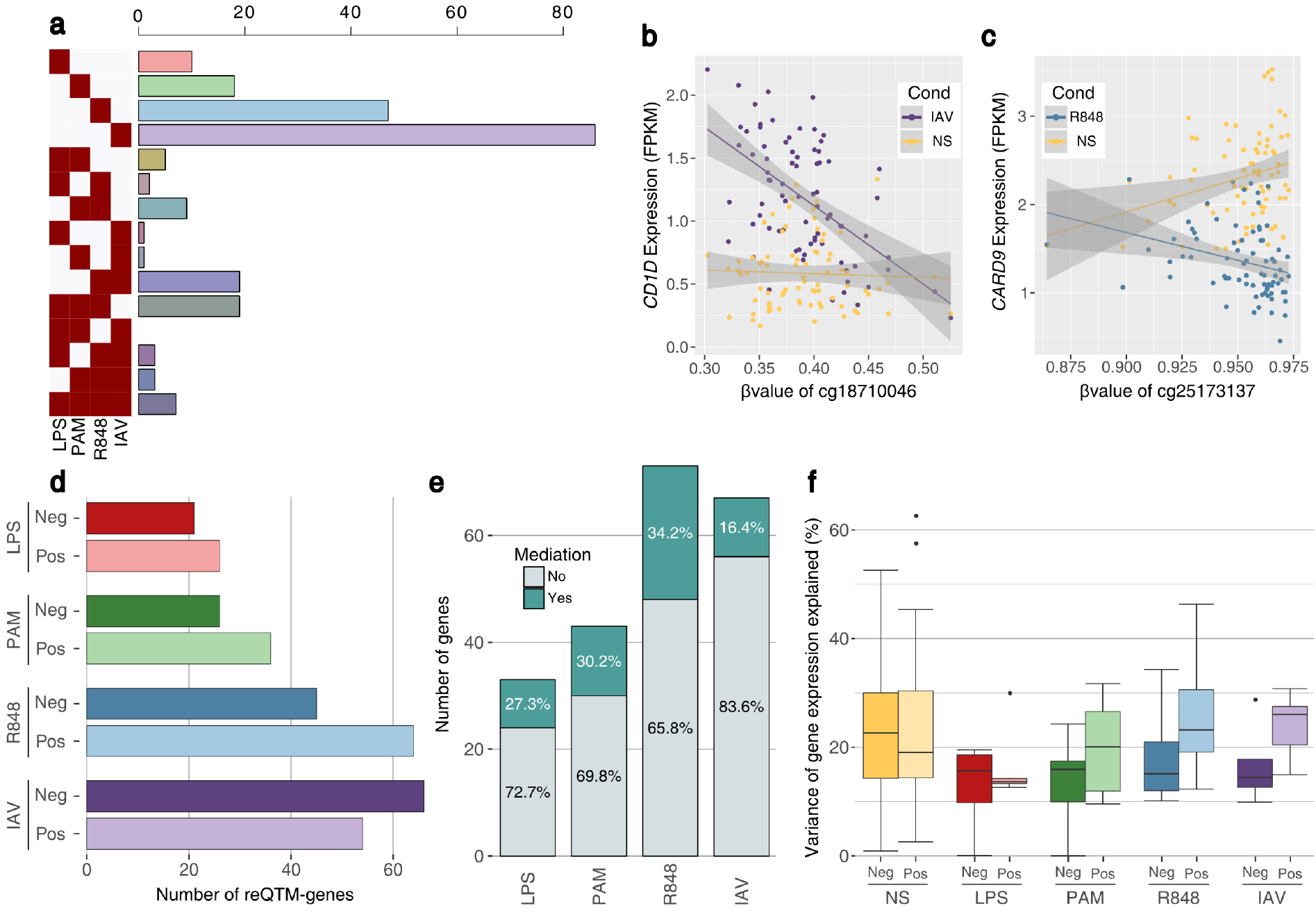
Effects of DNA methylation on transcriptional responses to immune stimulation. **a** Number of genes harbouring reQTMs in single conditions or combinations of stimulations. **b-c** Examples of reQTMs detected in this study. Lines indicate the fitted linear regression model, and grey shades the 95% confidence intervals of these models. **b** The distribution of the expression values of *CD1D* at the non-stimulated (yellow) and after IAV infection (purple) is plotted as a function of β-values, for AFB individuals only. **c** The distribution of the expression values of *CARD9* at the non-stimulated (yellow) and upon R848 stimulation (blue) is plotted as a function of β-values, for EUB individuals only. **d** Number of reQTM-genes by condition and according to the direction of their association with DNA methylation. **e** Number of mediated and non-mediated reQTM-genes per stimulation condition. The percentages of these two categories for each condition are also indicated. **f** Proportion of variance of gene expression explained by DNA methylation, among negative (dark colours) and positive (light colours) associations, in mediated cases.

To explore causal mediation effects of DNA methylation in the context of immune activation, we mapped response-QTLs (see **“Methods”**). Following our previous rationale (**Figure S9**), we identified 141 trios (61.3% of the 230 reQTM-genes, **Table S4**). At FDR=5%, we detected 40 genes (28.4%) where the genetic control of their transcriptional response was mediated by DNA methylation (**Fig. 5e**). Although non-significant, we found a higher proportion of mediation for genes whose response was positively associated with DNA methylation, as compared to negative associations, in particular for viral challenges (OR ~2.0; Fisher’s exact *P* = 0.33) (**Figure S14)**. Among mediated genes in the viral conditions, the proportion of gene expression variance explained by DNA methylation was higher for positive than for negative associations, again at odds with the non-stimulated condition (**Fig. 5f**). More generally, our analyses illustrate the value of mapping reQTMs and studying the underlying patterns of causality, to uncover mechanisms that might explain disparities in the way individuals and populations respond to immune activation.

## Discussion

Our population epigenetic results, obtained in the setting of an innate immunity cell population, demonstrate extensive differences in DNA methylation profiles between two populations that differ in their genetic ancestry but share the same present-day environment. Such population differences were observed at the epigenome-wide level (explaining ~12% of the total variance in DNA methylation) and involved 12,050 sites that were mostly located in genes with functions related to cell periphery or immune response regulation. A first interesting insight that can be drawn from these analyses is that genes involved in the activation and regulation of immune responses tend to present higher levels of DNA methylation in individuals of European ancestry, with respect to those of African-ancestry, mostly owing to genetic control. This intriguing observation could provide a mechanistic explanation for the ancestry-related differences in transcriptional responses to bacterial pathogens recently reported in macrophages, where European ancestry is associated with lower inflammatory responses [49].

We found that 70% of differentially methylated sites between populations were associated with at least one meQTL, supporting the notion that population differences in DNA methylation are mostly driven by DNA sequence variants [38, 40-42]. In some cases, a single genetic variant can account for important population differences at multiple CpG sites, as attested by the *trans*-meQTL we detected at *CTCF*, whose local genetic variation has been shown to alter distant DNA methylation patterns in whole blood [65]. We show that a *CTCF* variant (rs7203742) regulates DNA methylation of 30 distant CpGs, 40% of which are differentially methylated between populations. We also found that most *CTCF trans*-regulated CpGs are located nearby *CTCF* binding sites (mean distance 1,984 bp) but, interestingly, even closer to binding sites of other TFs (mean distance 44 bp, Wilcoxon *P =* 1.7×10^−9^) with 60% of them falling directly within the TF binding site. This observation is consistent with a model of pioneer transcription factor activity [94], and suggests that *CTCF* acts as a pioneer factor that will generate changes in chromatin state that, in turn, will become accessible for binding of secondary factors.

This study also establishes that inter-individual differences in DNA methylation are associated with gene expression variation (eQTMs) mostly following a canonical model of negative associations, particularly in enhancer regions. At the population level, we find that the extent of sharing of eQTMs between individuals of African and European ancestry is significantly higher than that of meQTLs (62% *vs*. 50%; Fisher’s exact *P* = 1.73×10^−15^). This suggests that the links between DNA methylation and gene expression are more stable across populations than the genetic control of DNA methylation itself, an observation that cannot be explained by differences in power between these analyses (**Supplementary Note 2**).

At the genome-wide level, we find that the quantitative impact of DNA methylation on gene expression variation is lower than reported by some previous studies, possibly reflecting differences in experimental settings and statistical power (e.g., cell types, sample sizes, etc.) [23, 65, 83, 88]. For example, a study of 204 healthy new-borns detected substantial variation across tissues in the number of genes whose expression levels were associated with DNA methylation, ranging from 596 in fibroblasts to 3,838 in T cells [23]. We detected, at the non-stimulated state, 811 eQTM-genes (6% of the total number of expressed genes), a figure that drops to 230 for reQTM-genes across stimulation conditions. However, a limitation of our study is that we measured DNA methylation at the basal state, while gene expression was obtained after 6 hours. Studies including a more comprehensive range of epigenetic marks obtained at different time points — in different cell types and tissues originating from individuals of various ancestries — are needed to more precisely understand the interplay between these regulatory elements and quantify their respective roles in the regulation of transcriptional activity.

The detected eQTMs were found to be drastically enriched in genetic control (OR ~33.2, *P* < 1×10^−326^, **Fig. 3c**), which highlights the coordinated action of genetic and epigenetic factors in driving gene expression variation but raises questions about the causal role of DNA methylation [56]. Despite cautious interpretation of causality in mediation analyses is required [95], our analysis provides a first estimate of the potential direct role of DNA methylation in regulating transcriptional activity, in both resting and stimulated monocytes. At the non-stimulated state, we find that ~20% of eQTM-genes show evidence of a causal mediation effect of DNA methylation. Although a similar extent of mediation was found upon immune stimulation (~17%), we detected specific patterns upon treatment with viral challenges, where a higher occurrence of positive associations was observed among mediated cases. These findings mostly reflected cases where high levels of DNA methylation were associated with low gene expression in the non-stimulated condition, thus requiring stronger responses to reach high levels of gene expression upon cell perturbation. These trends suggest a major, direct and context-specific role of DNA methylation in the regulation of immune responses, whose complexity requires further investigation.

Finally, we found that meQTLs, in particular those associated with ancestry-related differences, are enriched in GWAS hits related to immune disorders. This suggests that DNA methylation might have an important impact on the cellular activity of monocytes and ultimately affect phenotypic outcomes. Nonetheless, a large fraction of the variance of DNA methylation and gene expression remains unexplained. Additional work is needed to quantify the relative impact of genetic, epigenetic, environmental, and lifestyle factors in driving variation of DNA methylation and gene expression, both in resting and stimulated cells. Furthermore, although the causal mediation analyses presented in this study reinforce the notion that DNA methylation can play a direct role in regulating gene expression in humans [23, 96], monitoring the kinetics of variation in DNA methylation and gene expression after exposure to different infectious agents will broaden our understanding of the interplay between these molecular phenotypes and their impact on end-point phenotypes.

## Conclusion

Our study reveals extensive variation in DNA methylation profiles between individuals and populations, with ancestry-related differences being mostly explained by genetic variation. It also suggests that DNA methylation can have a direct, causal impact on the transcriptional activity of primary monocytes, providing new insight into the nature of the host factors that drive immune response variation in humans.

## Materials and Methods

### Sample collection and monocyte purification

The EvoImmunoPop collection consists of 200 individuals (males between 20-50 years old, mean: 31.5 years old) from two different ancestries (100 of European and 100 of African descent), who were recruited at the Center for Vaccinology from the Ghent University Hospital (Ghent, Belgium) [48]. For each participant, 300 ml of whole blood was collected into anticoagulant EDTA-blood collection tubes and peripheral blood mononuclear cells (PBMCs) were purified using Ficoll-paque density gradients (#17-1440-03, GE Healthcare). Monocytes were positively selected from purified PBMCs using magnetic CD14 microbeads (#130-050-201, MiltenyiBiotec), as per manufacturer’s instructions. All samples had a monocyte purity higher than 90% with a mean value of 97%.

### DNA Methylation profiling and data normalization

Genomic DNA was extracted from the monocyte fraction using a phenol/chloroform protocol followed by ethanol precipitation. The DNA was then bisulfite converted and BC-DNA was then processed using the Illumina Infinium MethylationEPIC BeadChip Kit (Illumina, San Diego, CA) to obtain the methylation profile of each individual at more than 850,000 CpG sites genome-wide.

In total, 184 samples were hybridized with the EPIC array, including 172 unique samples and 12 technical replicates. We removed any technically unreliable probes: (i) potentially cross-hybridizing probes, (ii) those located on the X and Y chromosomes, as well as (iii) probes overlapping SNPs that present a frequency higher than 1% in at least one of the studied populations. These SNPs were chosen based on our own genotyping dataset, as well as on the 1,000 Genomes project [97]. To control for the quality of the probes and samples, we filtered out individuals with > 5% of probes associated with a detection *P*-value > 10^−3^, and then, probes with a detection *P*-value > 10^−3^ in one or more individuals. Following this filtering process, 552,141 of the original 866,837 sites on the array were retained.

We calculated methylation levels from raw data, using the R Bioconductor lumi package [98]. Given that the M-value has been shown to provide better detection sensitivity than β-values at extreme levels of modification [68], we used the M-value to run all statistical analysis unless otherwise stated. Note that in some instances of the text and figures, β-values are reported for ease of clarity and interpretation. M-values were then adjusted for background noise with the Normal-exponential using out-of-band probes (noob) from the R Bioconductor minfi package [99]. Next, normalization for colour bias was performed using *lumiMethyC* with the ‘quantile’ method, and for methylated/unmethylated intensity variation using the *lumiMethyN* with the ‘ssn’ method [98]. Finally, we corrected for technical differences between type I and type II assay designs, by performing Beta-mixture quantile normalization [100]. To correct for known batch effects and potential hidden confounders, we used the *sva* function from the sva Bioconductor package [101] with age as a variable of interest. Additionally, five EUB samples were removed because they presented an excess of hemimethylated sites, leaving 89 EUB and 78 AFB samples. To obtain equal power in the two studied populations, we down-sampled the European group to 78 samples by randomly removing 11 EUB samples, for an overall final cohort of 156 individuals.

### Extraction of differentially methylated sites (DMS)

To detect CpG sites presenting statistically different levels of DNA methylation between AFB and EUB, we fitted a linear regression model for each CpG site: M-value ~ population + age + surrogate variables + error, and next applied an empirical Bayes smoothing to the standard errors using the R Bioconductor limma pipeline [102]. *P*-values were adjusted using the Benjamini & Hochberg method. DMS were extracted using a threshold of adjusted *P*-value (<0.01) and a difference in the mean β-value of each population |*∆*β| > 5%.

### Mapping of methylation quantitative trait loci (meQTLs)

All individuals were genotyped for a total of 4,301,332 SNPs on the Illumina HumanOmni5-Quad BeadChips, and went through whole-exome sequencing with the Nextera Rapid Capture Expanded Exome kit, on the Illumina HiSeq 2000 platform, with 100-bp paired-end reads. Details of the processing of genotyping and whole-exome sequencing data, together with imputation using the 1,000 Genomes Project imputation panel [97], are reported in ref. [48]. For the meQTL mapping, we filtered out SNPs with a minor allele frequency < 5% in the populations studied, and kept a final dataset of 10,278,745 SNPs (i.e., corresponding to the merged genotyping and whole-exome sequencing dataset after imputation; 8,913,090 SNPs in Africans and 6,178,808 SNPs in Europeans). Age, PC1 and PC2 of the genotype matrix, and surrogate variables were used as covariates in the linear model.

We mapped meQTLs using the statistical framework implemented in the MatrixEQTL R package [69]. For local associations (*i.e.*, distance SNP-CpG ≤ 100kb), we performed two independent mappings using (i) the direct linear model from the MatrixEQTL pipeline, and (ii) a Kruskal-Wallis rank test. Associations were considered significant when passing the 5% FDR threshold in both mappings. Two models were considered: merging all individuals and including a binary variable adjusting for ancestry or keeping the two populations separately. To detect all possible independent SNPs regulating methylation at a single CpG site in *cis*, we regressed out genotypes of all primary *cis*-meQTLs and then performed *cis*-meQTL mapping on the regressed methylation data to find secondary *cis*-meQTLs. We repeated this process in a stepwise fashion until no additional independent *cis*-meQTLs were detected. This allowed us to refine our local meQTL mapping by detecting all possible independent SNP-CpG associations.

For distant, *trans*-acting associations (*i.e.*, distance between SNP and CpG ≥ 1Mb or on different chromosomes), we restricted our analysis to SNPs located in the vicinity of transcription factor (TF) coding genes, to limit the burden of multiple testing. We selected all SNPs located less than 10kb to the TSS of any expressed TF in our dataset. For each SNP, we only investigated CpG sites that mapped at least 1 Mb from the SNP or located on other chromosomes, using a Kruskal-Wallis rank test.

For both *cis*- and *trans*-meQTLs, FDR was computed by mapping meQTLs on 100 datasets with the M-values permuted within each population. We then kept, after each permutation, the most significant *P*-value per CpG site, across populations (probe-level FDR). Finally, we computed the FDR associated with different *P*-value thresholds for *cis* or *trans*, and subsequently selected the *P*-value threshold that provided a 5% FDR: *P* = 1×10^−5^ and *P* = 1×10^−9^ for *cis*- and *trans*-meQTLs, respectively.

### Investigating the genetic basis of population differences in DNA methylation

We aimed at identifying the proportion of the population differences in DNA methylation that was accounted for by genetic variability. To do so, for the 8,459 DMS that were associated with at least one meQTL, we computed the following ratio:

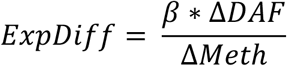

with β reflecting the effect of the derived allele of the meQTL on methylation, ∆DAF the difference in allelic frequencies between Europeans and Africans (DAF_EUB_ – DAF_AFB_), and ∆Meth the observed difference in the mean levels of DNA methylation between European and African individuals (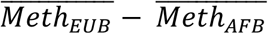).

Note that this ratio is not bound to [0:1], as the effect of genetics onto the overall population differences in DNA methylation can be counteracted by opposite effects of independent origins (e.g. environmental factors or non-detected independent genetic effects).

### Detecting population-specific meQTLs

We aimed at distinguishing population-specific meQTLs (i.e. SNPs present at similar frequencies in both populations but having different effect-sizes on DNA methylation between populations) from meQTLs detected in one population only due to population differences in allelic frequencies. We considered as population-specific, meQTLs whose effect size was significantly different between populations. To do so, we fit the following linear model:

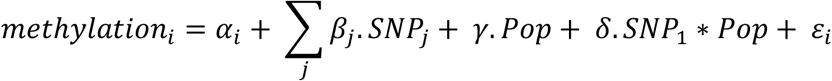

where SNP_j_ is the genotype of the j-th variant, Pop is a binary variable indicating the population origin (0 for Europeans and 1 for Africans), and ε_i_ is a random, normally distributed residual. In this model, the β_j_ reflects the effect of the derived allele of the SNP_j_ on methylation, γ estimates the fold change in methylation between populations observed for individuals with identical genotype, and δ captures the differences in the primary meQTL effect size between populations. Such a model allows to test for a difference in meQTL effect size between populations by testing the null hypothesis, δ = 0 (interaction test). We considered meQTLs as being population-specific when the adjusted interaction *P*-value at the locus was lower than 0.05 (corresponding to FDR < 5%).

### GWAS enrichment analyses

We used the NHGRI GWAS catalog [103] to first select all significant SNPs that were significantly associated with a complex trait or disease at a *P* < 1×10^−8^. Using this set of GWAS hits, we next extracted all SNPs in LD with each of these hits (R^2^>0.8), and classified the resulting final set of 166,248 SNPs according to their parental Experimental Factor Ontology (EFO) term [74].

We then selected all meQTLs in our dataset that passed the *P*-value threshold corresponding to FDR 5% in our initial mapping, and filtered out meQTLs that were in LD (R^2^>0.8) keeping one SNP per independent loci (56,574 independent SNPs). For the resampling set, we considered all SNPs that were initially used for the meQTL mapping and pruned them for LD (R^2^>0.8), yielding a final set of 921,466 SNPs. Resampling was performed using bins of allelic frequencies at intervals of 5%.

Finally, we tested for fold-enrichments of meQTLs in GWAS hits, for each of the 17 parental EFO categories [74]. The fold-enrichment was calculated by comparing the number of LD pruned-meQTLs that were found to correspond to GWAS hits (or were in LD with GWAS hits) with the expected number estimated through 10,000 resamples. *P*-values associated to the fold-enrichment were calculated by fitting a normal distribution to the empirical distribution of our 10,000 resampled sets of SNPs. Confidence intervals were computed using 10,000 resamples by bootstrap. The same procedure was applied when searching for enrichments of meQTLs specifically in GWAS hits related to the 268 traits of the “Immune system disorder” EFO parental term.

### Expression quantitative trait methylation (eQTM) analysis

To identify associations between DNA methylation levels and gene expression of nearby genes, we leveraged RNA-sequencing data obtained from the same individuals, both at the non-stimulated state (NS) and in response to four immune stimuli [48]. Briefly, RNA-sequencing was performed on the Illumina HiSeq2000 platform with 101-bp single-read sequencing with fragment size of around 295 bp, and outputs of around 30 million single-end reads per sample were obtained. A total of 763 RNA-sequencing samples from our filtered dataset of 156 donors were analysed for gene expression profiling, including 156, 151, 153, 148 and 155 samples for the NS, LPS, Pam3CSK4, R848 and IAV conditions, respectively. Details of cell culture, immune stimulation conditions, and RNA-seq processing can be found in ref. [48].

Using the RNA-sequencing data from the NS condition, we mapped eQTMs (i.e. CpGs whose variation is associated with gene expression) in a window of 100 kb around the TSS of each gene (12,578 expressed genes in primary monocytes). The associated *P*-values and the coefficients of correlation between methylation profiles and gene expression were obtained using a Spearman’s rank correlation. FDR was computed by mapping eQTMs on 100 datasets with the M-values permuted, and kept, after each permutation, the most significant *P*-value per gene (gene-level FDR). We selected the *P*-value threshold that provided a 5% FDR (*P* = 5×10^−5^).

We also mapped eQTMs in the context of the response to the various stimulations, namely response-QTMs (reQTMs). To do so, the same procedure explained above for the eQTM mapping was followed, using the fold-change of expression upon stimulation as a measure of the host response to infection. Specifically, we calculated the difference of the log_2_ of expression values between the stimulated and non-stimulated states, corrected for the effect of low-values of FPKM, for each gene expressed in at least one of the two conditions.

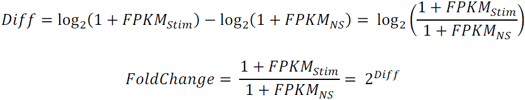

For the mapping of eQTMs and reQTMs, we conducted two separate analyses: merging all individuals and including ancestry as a covariate, or keeping the two populations separately.

### Expression quantitative trait loci (eQTL) analysis

We mapped expression quantitative trait loci (eQTLs) using the MatrixEQTL R package [69], leveraging our genotyping and expression data [48]. As for the meQTL mapping, we filtered out SNPs with a minor allele frequency < 5% in the populations studied and kept a final dataset of 10,278,745 SNPs. Age and PC1/PC2 of the genotype matrix were used as covariates in the linear model. Two different models were used: merging all individuals and including ancestry as a covariate, or keeping the two populations separately. We also mapped response quantitative trait loci (reQTLs), using the fold-change of expression described above, instead of expression, and the same covariates that we used for the eQTL mapping.

For both eQTLs and reQTLs, FDR was computed by mapping eQTLs/reQTLs on 100 datasets with the expression values permuted within each population. We then kept, after each permutation, the most significant *P*-value per gene, across populations (gene-level FDR). Finally, we computed the FDR associated with different *P*-value thresholds for eQTLs or reQTLs, and subsequently selected the *P*-value threshold that provided a 5% FDR: *P* = 5×10^−5^ and *P* = 5×10^−6^ for eQTLs and reQTLs, respectively.

### Simulations to infer causality

We simulated different scenarios to infer causal relationships between DNA methylation and gene expression. For each scenario, we started by randomly selecting genomic blocks of 1 Mb each along the genome to keep realistic expectations of genetic structure. We next randomly sampled SNPs in these blocks, which we used to simulate methylation and gene expression data. For example, in a scenario where a genetic variant influences DNA methylation variation that, in turn, actively regulates gene expression (see **Fig. 4a**), we followed the next steps:

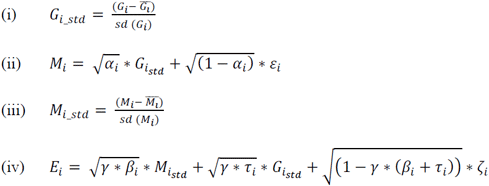

where G_i_ is the genotype of the i-th sampled variant and G_i_std_ the standardized value of its genotype; M_i_ is the simulated methylation data and M_i_std_ its standardized methylation value; E_i_ is the simulated gene expression data; *α*_i_, is the proportion of variance of M_i_ that is explained by G_i_, and γ is a noise parameter that corresponds to the total proportion of variance of E_i_ that is explained by G_i_ and M_i_. β_i_ and τ_i_ are the proportions of explained variance that are attributable to G_i_ and M_i_ respectively (satisfying β_i_+τ_i_=1). Finally, ε_i_ and ζ_i_ are random, normally distributed residuals. Note that in the simulation presented in **Fig. 4a-b**, we used a gamma of 0.25, so that 75% of the variance of gene expression remained unexplained.

### Detection of genetic variants-DNA methylation-gene expression trios

To infer causality between regulatory loci and gene expression variation, we considered eQTLs that were also detected as meQTLs, and, out of this subset, we kept only those for which the meQTL-CpG had previously been identified as an eQTM of the eQTL-gene (see **Figure S9** for clarity). When multiples SNPs or CpGs where present in a trio, we used an elastic net model, to build linear predictors of (i) gene expression based on DNA methylation variability for trios with multiple CpGs, and (ii) DNA methylation based on genetic variability for trios with multiple SNPs. These predictors were then used as summary variables for DNA methylation variability (i) or genetic variability (ii). Specifically, the *glmnet* function from the R package glmnet [104] was used to fit the generalized linear model via penalized maximum likelihood, with an elastic net mixing parameter α of 0.5. The strength of the penalty λ_1se_ was chosen as the largest value of lambda such that the error was within 1 standard deviation of the minimum lambda, when performing k-fold cross validation with the *cv.glmnet* function. Finally, the generic R function *predict* was used to build the optimal linear predictor in each case. For the trios presenting more than one SNP, we also used a predictor of gene expression based on genetic variability, as summary variable for the genetic variability, and found no differences in our simulation-based mediation results when compared to building the summary variable from a predictor of DNA methylation (data not shown).

### Mediation Analyses

For conducting causal mediation analyses, we used a Bayesian approach as implemented in the mediation R package [89]. Briefly, this approach estimates causal effects of a mediating variable *M* (DNA methylation) on the relationship between an independent variable *X* (genetics) and a dependent variable *Y* (gene expression). In this scenario, the global effect of *X* on *Y* can be written as *ρ_X→Y_* = *τ* + *α* ∙ *β*, where τ is the specific effect of *X* on *Y*, α the specific effect of *X* on *M*, and β the specific effect of *M* on *Y*. With this, the product α∙β represents the mediation effect of *G* on *Y*, through *M*. The *mediate* function of the mediation R package was used to compute point estimates for average causal mediation effects, as well as 1,000 simulation draws of average causal mediation effects. The empirical distribution of simulated effects was used to fit a normal distribution, which was subsequently used to compute empirical *P*-values for the H_0_ hypothesis “α∙β = 0”. We used the R function *p.adjust* with method “fdr” to correct at a FDR = 5%.

For comparison purposes with the mediation analyses, we conducted on simulated data a partial correlation approach to test for independence between expression and methylation levels when accounting for genetic variability. We used the *pcor.test* function from the R package ppcor [105] to compute *P*-values of the partial correlation between simulated expression and methylation data.

## Availability of data and materials

The DNA methylation data generated in this study have been deposited in the NCBI Gene Expression Omnibus (GEO) under accession code GSEXXXXXXXX. Genome-wide SNP genotyping, whole exome sequencing and RNA-sequencing data used in this study are available at the European Genome-Phenome Archive (EGA) under accession code EGAS00001001895.

## Authors’ contributions

L.T.H. designed and performed the computational analyses, analysed the data and interpreted the results, with input from M.R., M.F., H.Q., H.A., E.P. and L.Q.-M. L.M.M., J.L.M and M.S.K. contributed DNA methylation data. N.Z. contributed flow cytometry data. M.R., H.A. and E.P. contributed with ideas and participated in evaluating results and discussions. L.Q.-M. conceived and supervised the study and obtained the funding. L.T.H. and L.Q.-M. wrote the manuscript, with input from all authors. All authors approved the final manuscript.

## Ethics approval

All experiments involving human primary monocytes from healthy volunteers were approved by the Ethics Board of Institut Pasteur (EVOIMMUNOPOP-281297) and the relevant French authorities (CPP, CCITRS and CNIL).

## Additional File 1

**Figure S1.**
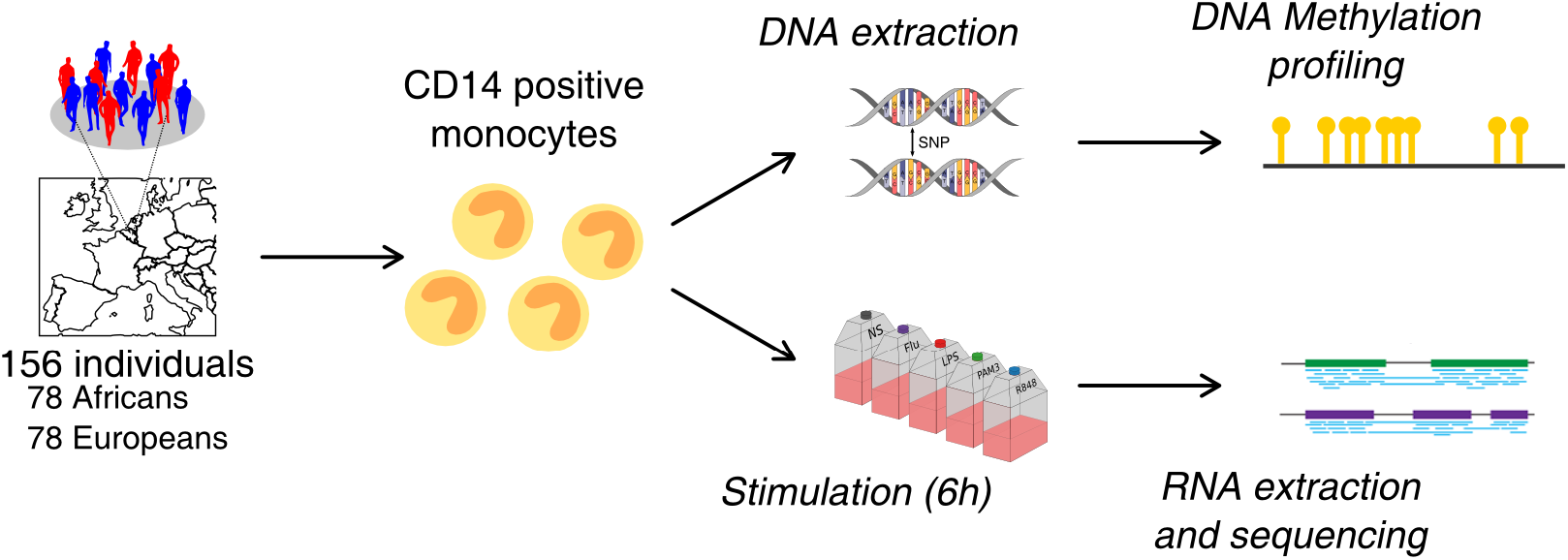
Overview of the EvoImmunoPop experimental setting. DNA methylation profiles and transcriptional responses to various immune stimulations, of primary monocytes from 156 healthy donors of European and African descent.

**Figure S2.**
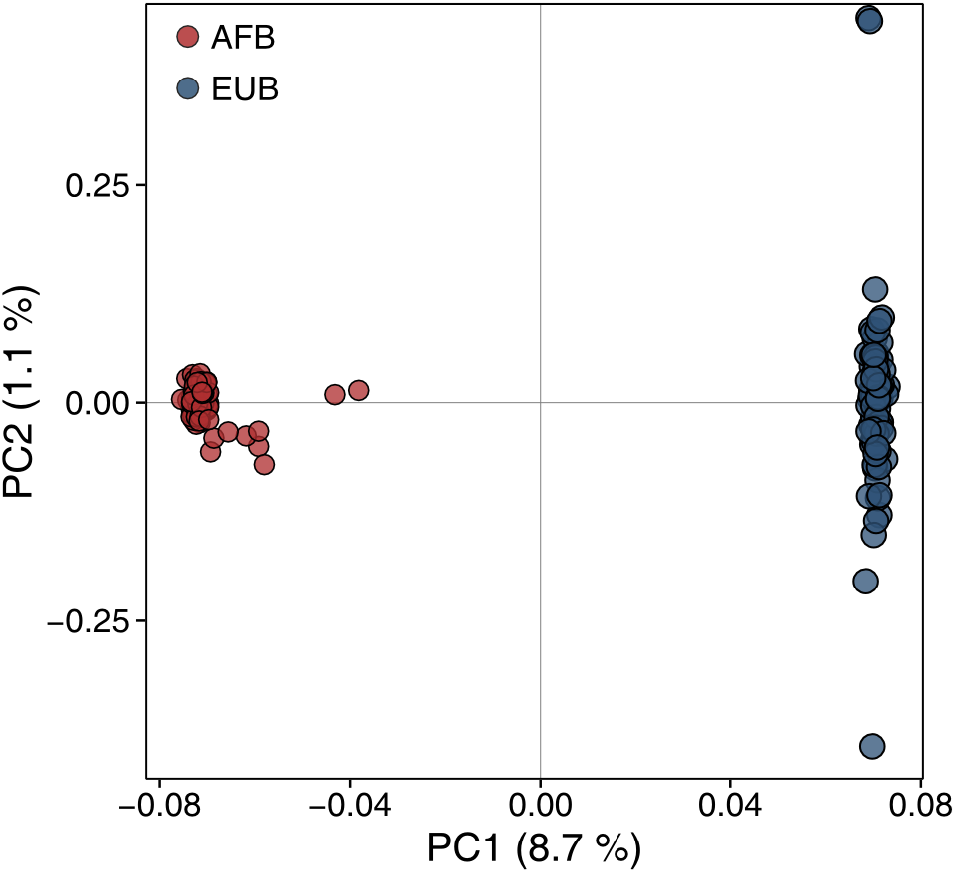
PCA of the genetic data, based on 151,419 SNPs, for Africans (AFB, red dots) and Europeans (EUB, blue dots). The percentages of variance explained by PC1 and PC2 are indicated.

**Figure S3.**
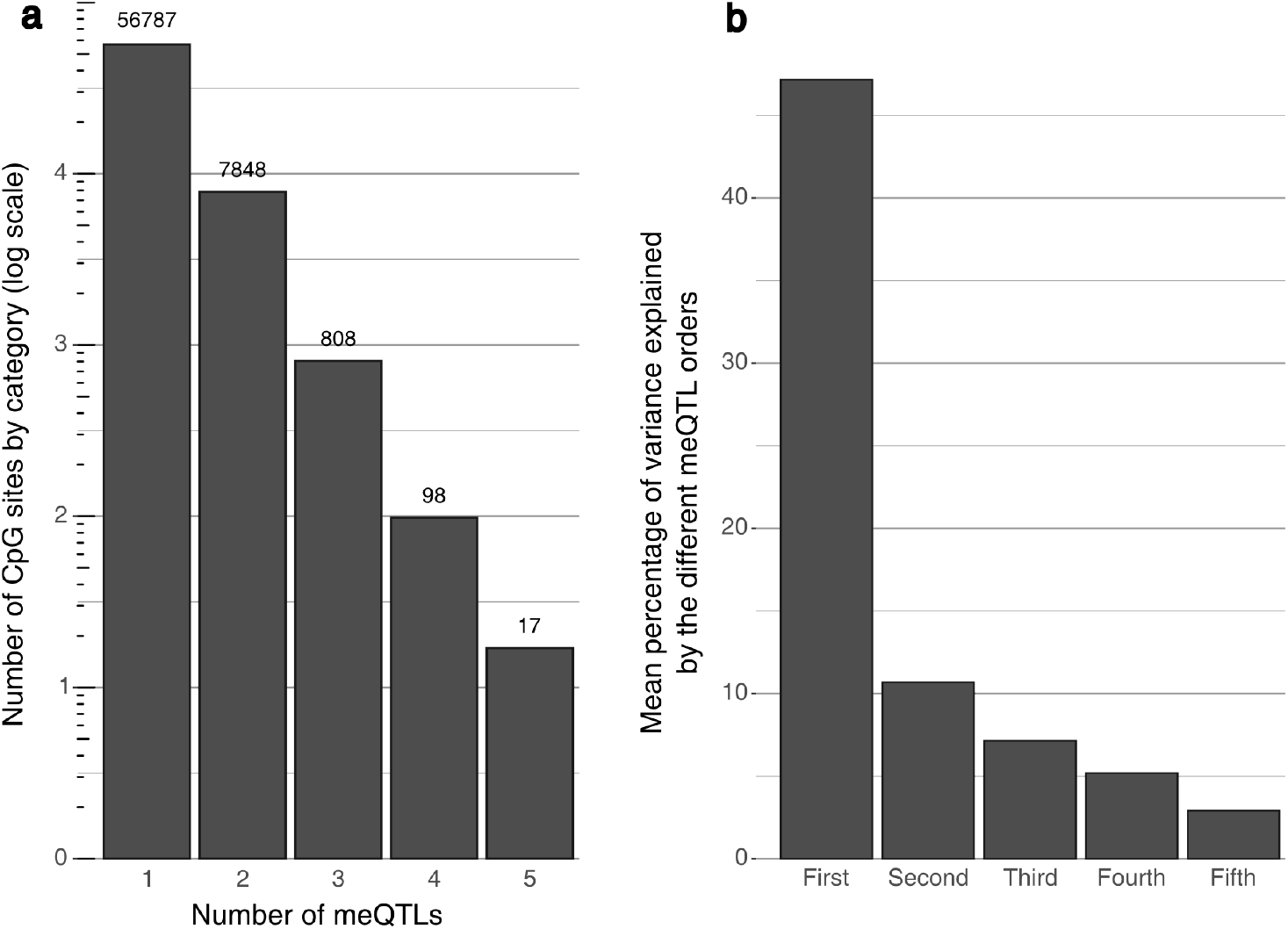
Fine mapping of meQTLs. **a** The number of CpG sites according to the number of associated independent meQTLs is shown on a log-scale. **b** Mean percentage of variance of DNA methylation explained by meQTLs according to the order in which they were detected.

**Figure S4.**
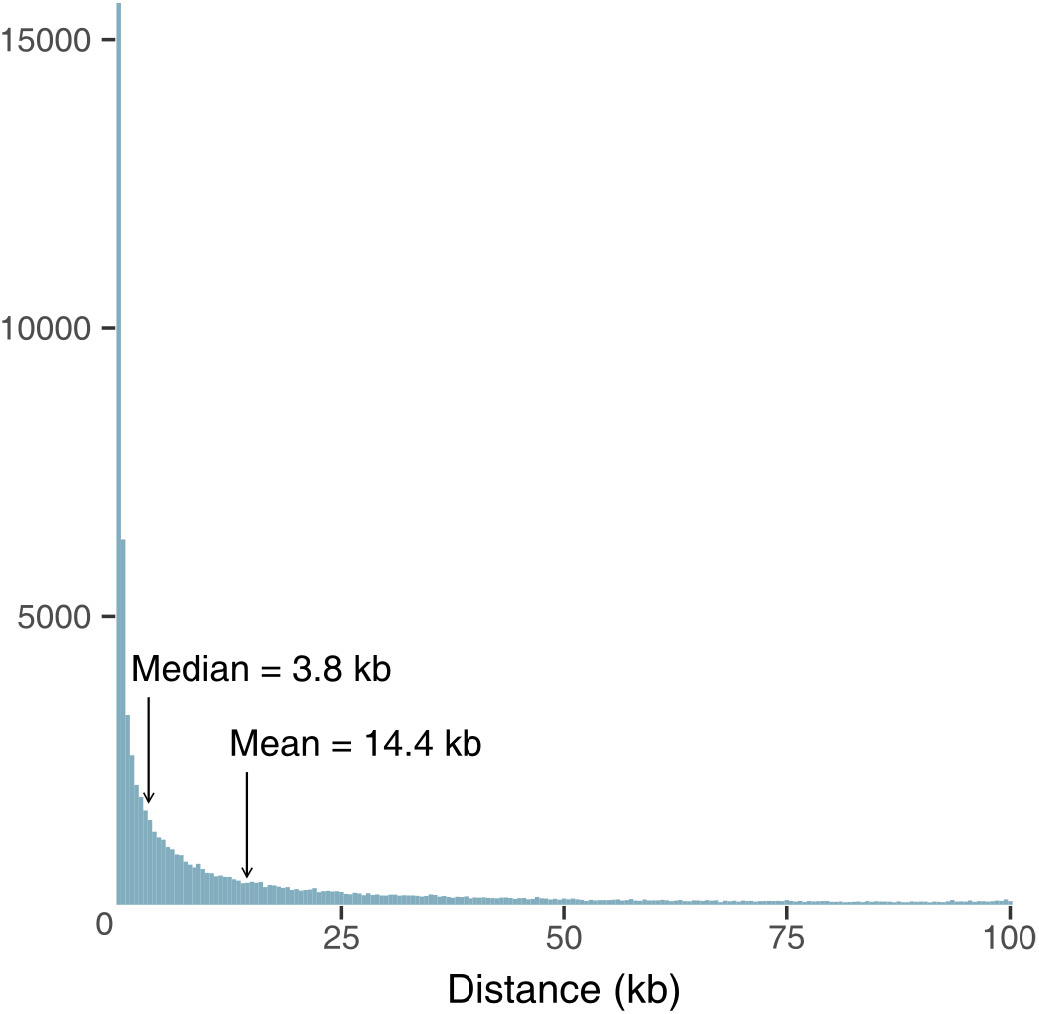
Histogram of physical proximity of *cis*-meQTLs. The distribution of the distances (in kb) between each meQTL and its associated CpG sites is presented, together with the mean and the median value.

**Figure S5.**
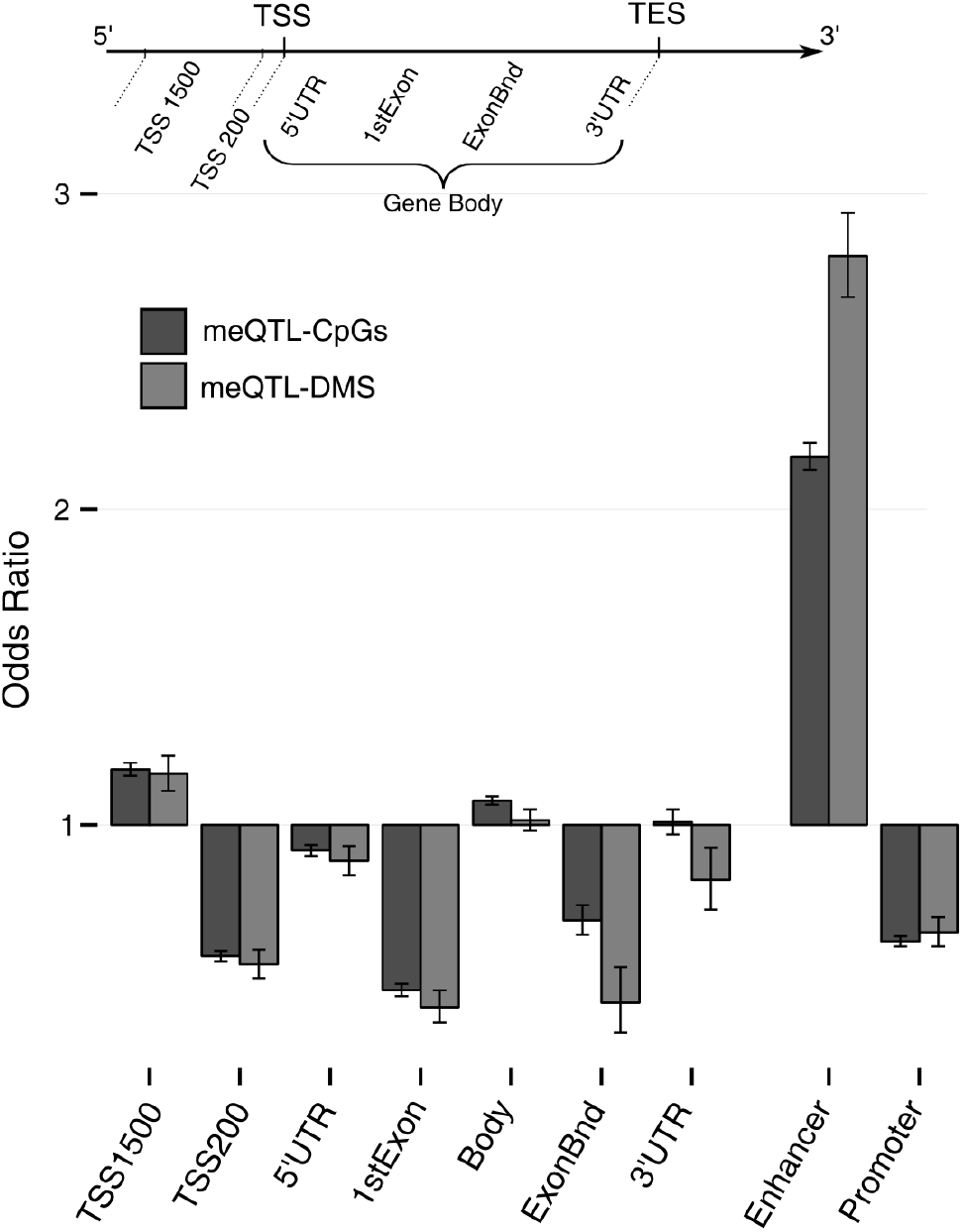
Genomic location of CpG sites associated with a meQTL. meQTL-CpGs are represented in dark grey, and the subset of these CpG sites that were also detected as DMS (meQTL-DMS) in light grey. OR were computed against the general distribution of the 552,141 CpGs of our dataset

**Figure S6.**
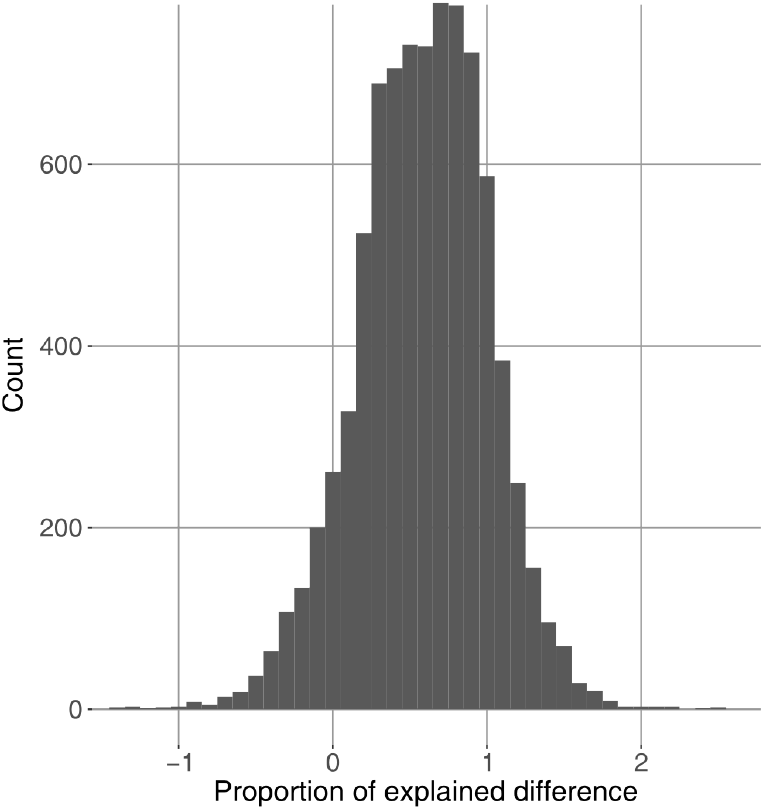
Proportions of population differences in DNA methylation accounted for by genetics. Histogram of the distribution of these proportions, for the 8,459 DMS that were associated with at least one meQTL. Proportions lower than 0 represent situations where genetics has an opposite effect to the observed overall population difference in DNA methylation. Conversely, proportions higher than 1 represent situations where the difference attributable to genetics is higher than that actually observed, indicative of an opposite effect of environmental factors or non-detected independent genetic effects.

**Figure S7.**
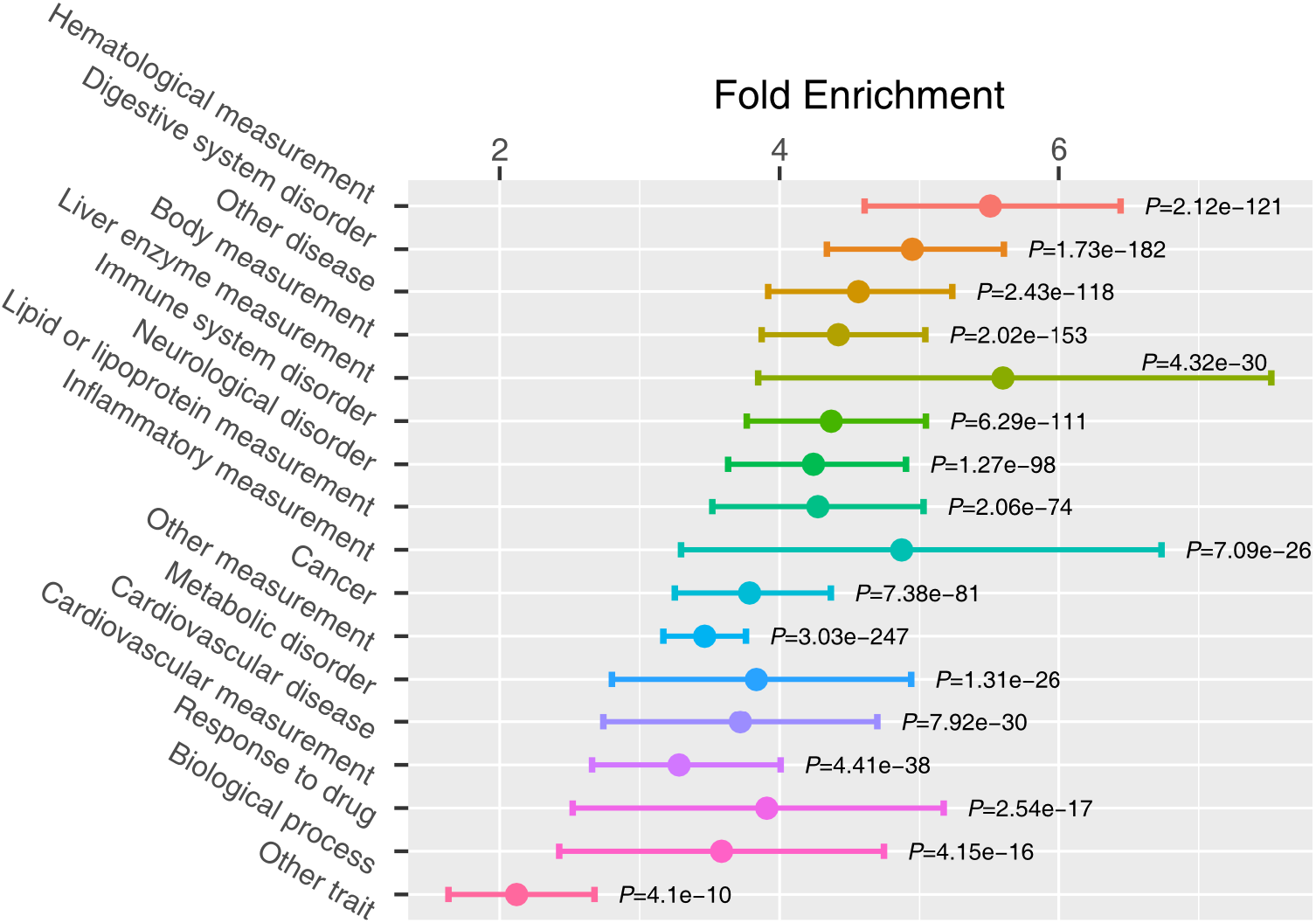
Fold enrichment of meQTLs in GWAS hits. For each of the 17 parental EFO categories, the fold enrichment, the 95% confidence intervals and the associated *P* values are shown.

**Figure S8.**
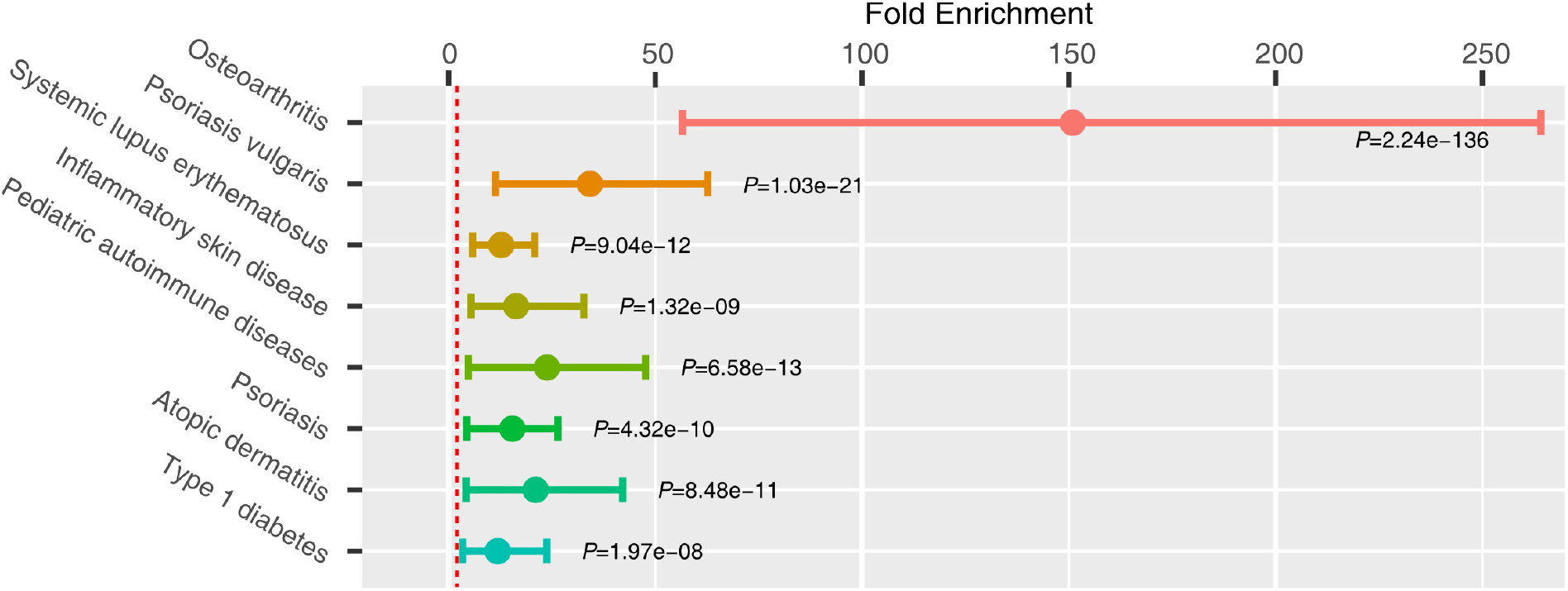
Fold enrichment of meQTLs associated with DMS in GWAS hits related to “immune system disorder”. For the 8 signals that presented the higher lower-bound of confidence intervals, the fold enrichment, the 95% confidence intervals and the associated *P* values are shown.

**Figure S9.**
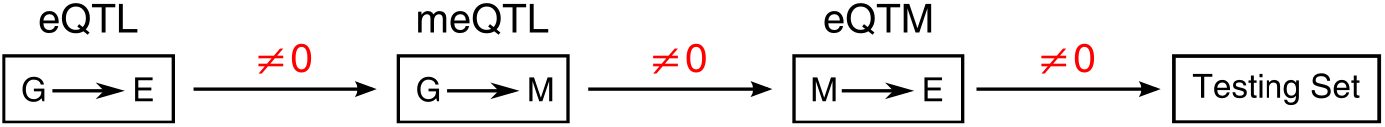
Rationale for the detection of trios to be used for causality inference.

**Figure S10.**
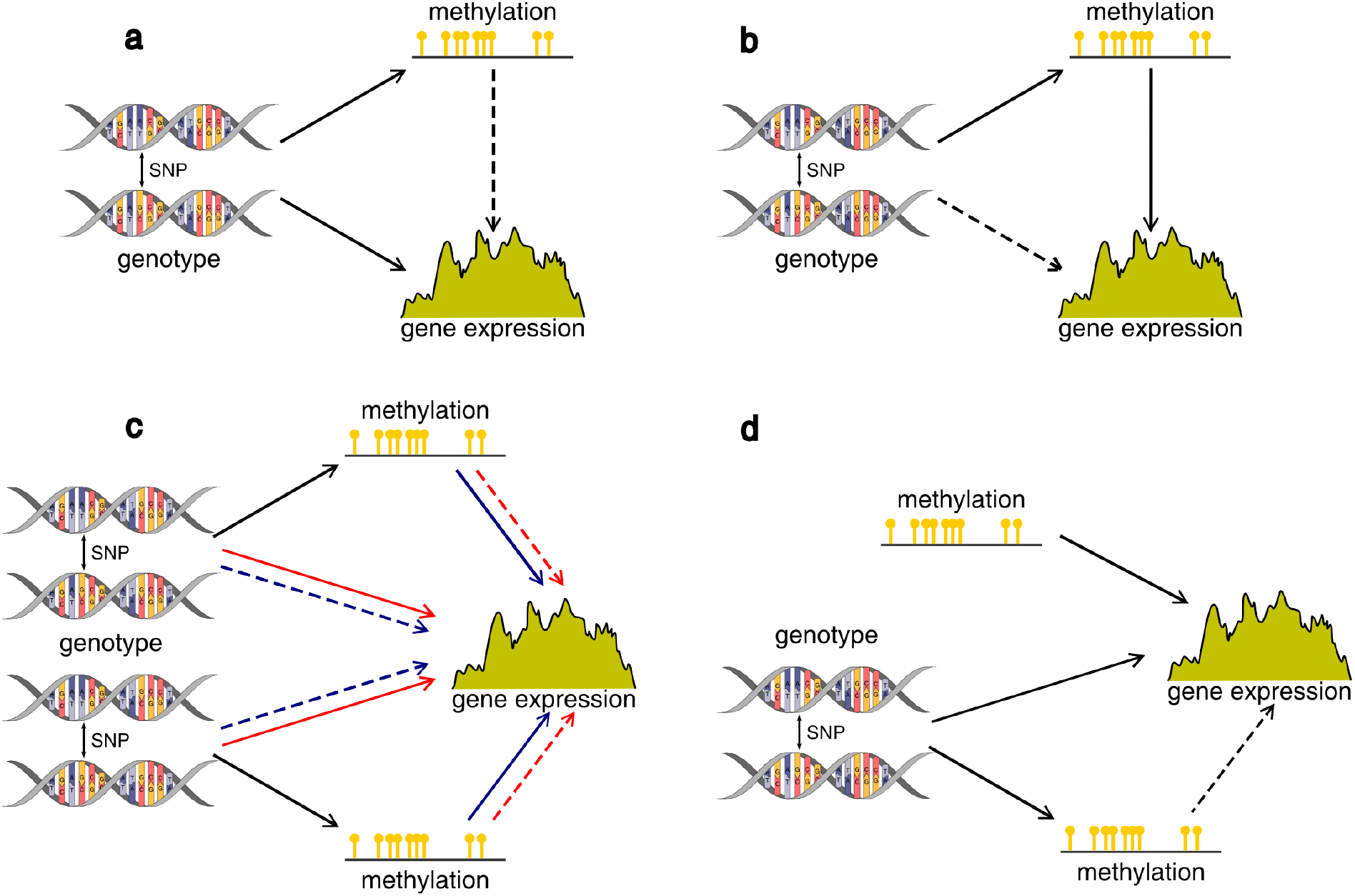
Cartoons of the various simulated scenarios. Plain arrows represent causal relationships, while dashed arrows represent correlations through another relationship. **a-b** Simple situations where either DNA methylation or genetics causally impact gene expression variation. **c** More complex scenarios where gene expression is causally impacted by two independent genetic (red arrows) or epigenetic (blue arrows) variants. **d** Scenario where the CpG site that causally impacts gene expression variation is not under the control of any genetic variant. Note that for all simulated scenarios (**a-d**), similar results between mediation analyses and partial correlations were obtained in terms of sensitivity and specificity (data not shown).

**Figure S11.**
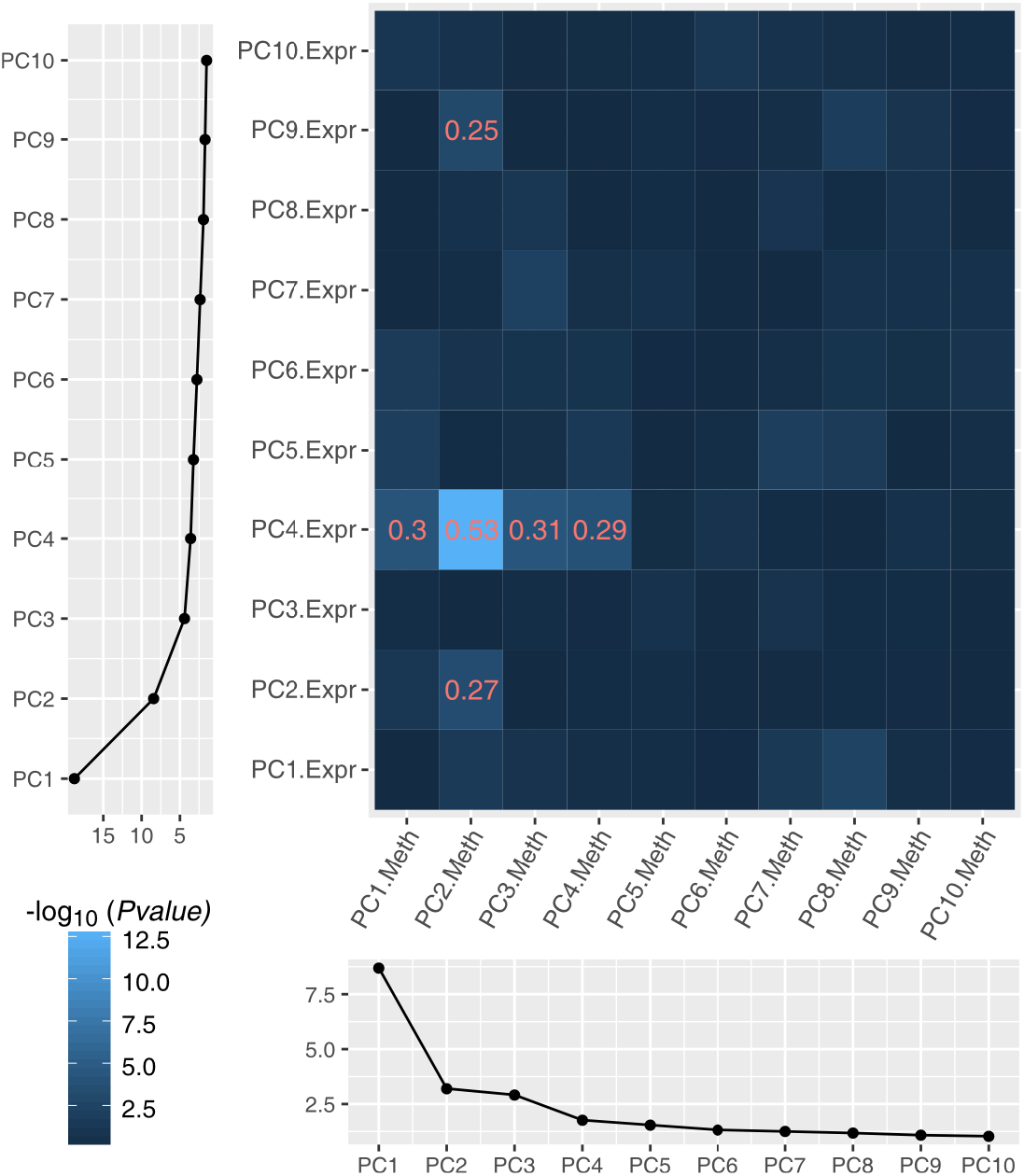
Heat map of correlation between the first ten PCs of expression and DNA methylation. Shades of blue are proportional with the −log10 of the correlation *P* values. In red are given the R^2^ of the correlation for cases were *P* < 0.001. Bottom and left panels show the percentage of variance explained by the first ten PCs of gene expression and DNA methylation, respectively.

**Figure S12.**
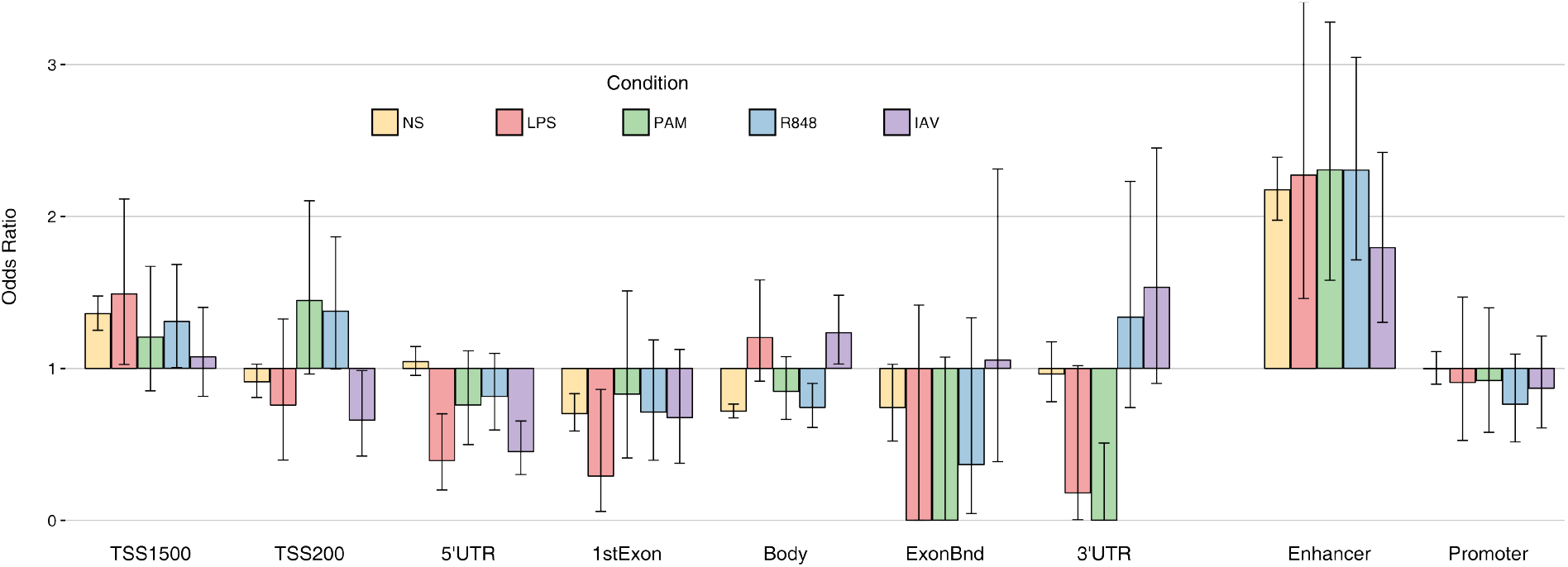
Genomic location of eQTMs (NS) and reQTMs (for all stimulated conditions). Odds ratio were computed against the general distribution of 552,141 CpGs of our dataset.

**Figure S13.**
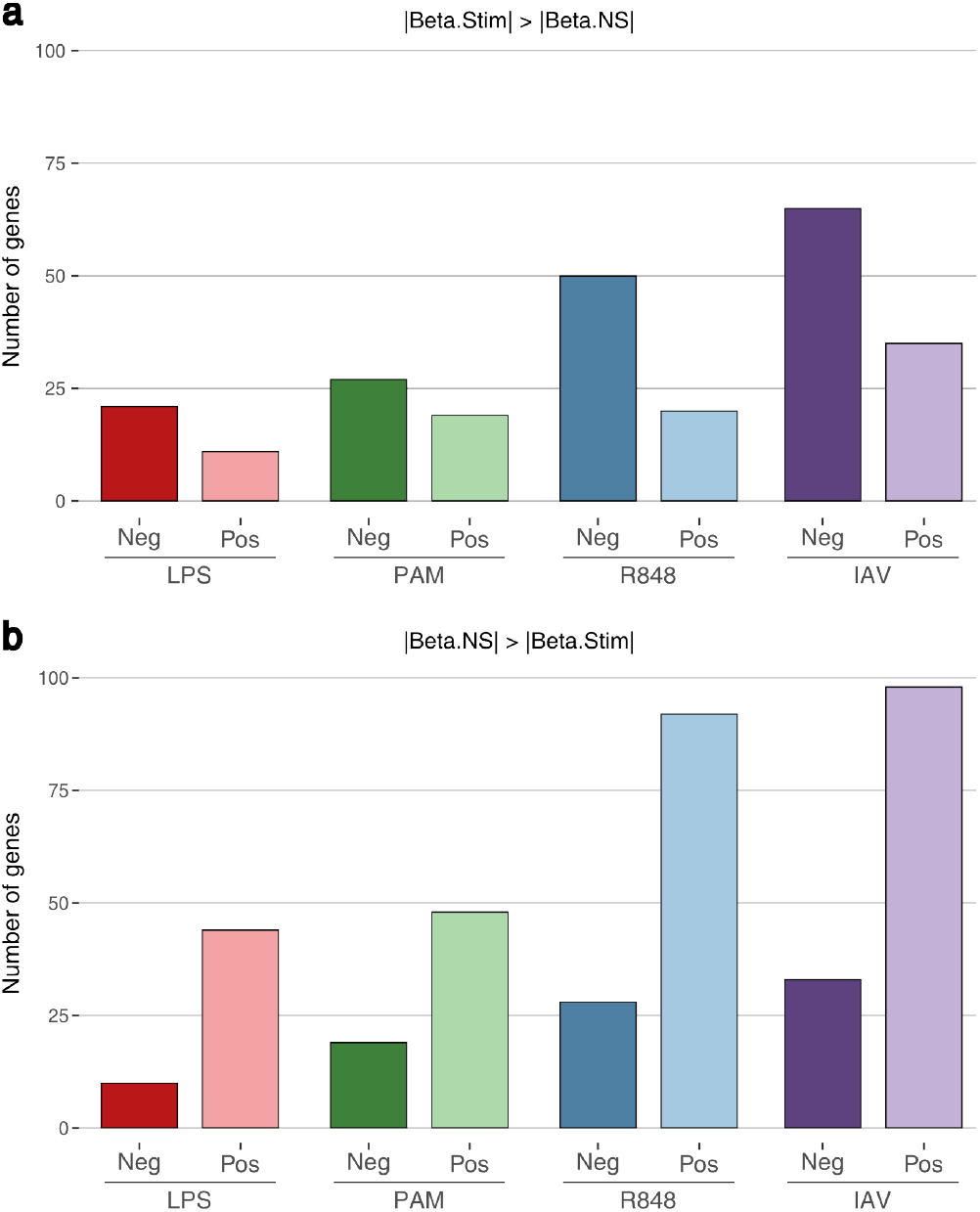
Number of reQTM-genes, per condition, according to the direction of their association with DNA methylation. **a** Cases presenting a stronger expression-methylation association upon stimulation than at the non-stimulated state, **b** Cases presenting a stronger expression-methylation association at the non-stimulated state than upon stimulation.

**Figure S14.**
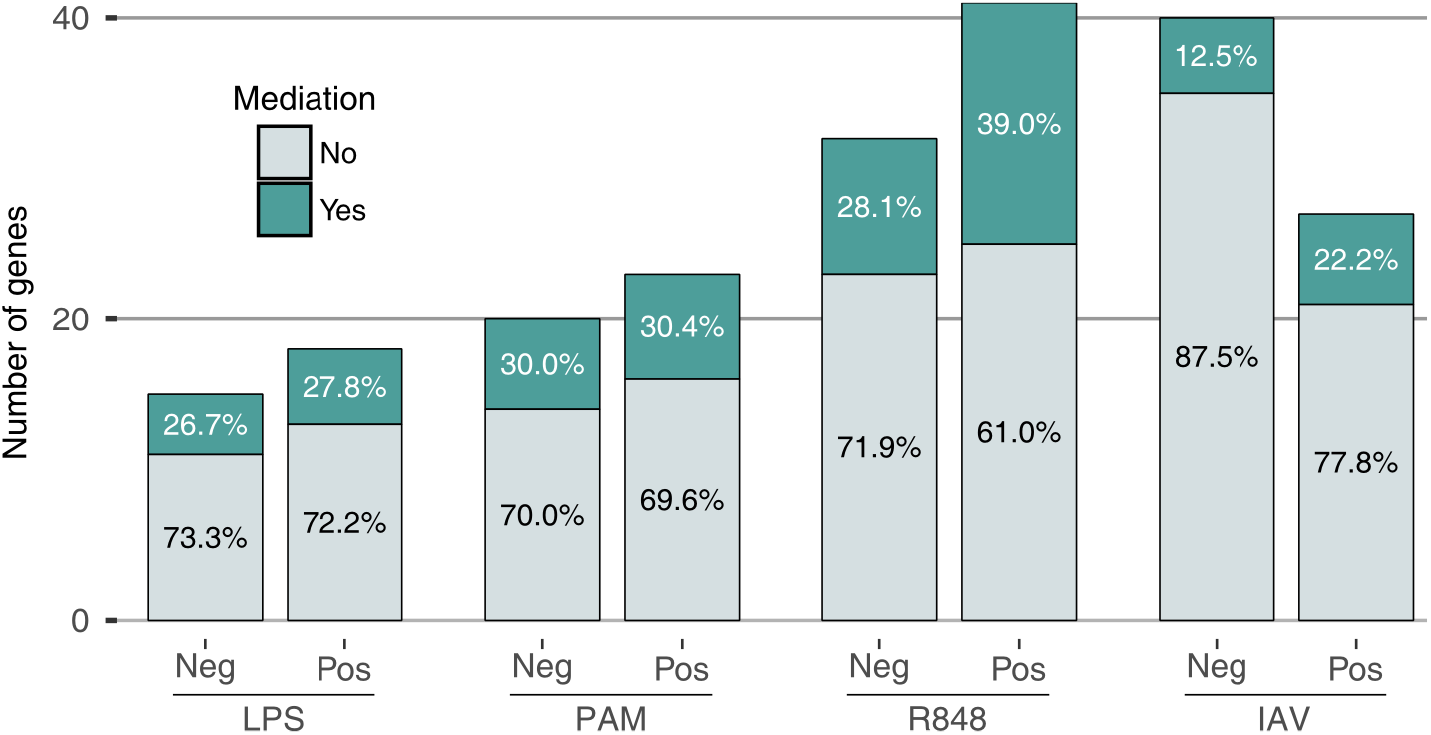
Causality inference upon immune stimulation. Number of mediated and non-mediated reQTM-genes for negative (Neg) and positive (Pos) associations between DNA methylation and fold-changes in expression upon different stimulation conditions. The percentages among these two categories are also indicated.

### Supplementary Note 1

The reverse causation scenario, where the impact of genetic variation on DNA methylation is mediated by gene expression variation, is highly unlikely in our experimental setting (**Figure S1**). Given that DNA methylation was obtained from monocytes at t=0, while gene expression was obtained after 6h, the reverse causation could only be observed in cases where expression at t=6h is a proxy of expression at t=0. We nonetheless tested this hypothesis by considering three different models: *Model 1*, independent control of both gene expression and DNA methylation by genetics; *Model 2*, genetic control of DNA methylation mediated by gene expression; and *Model 3*, genetic control of gene expression mediated by DNA methylation. We computed the log-likelihood of these three models:

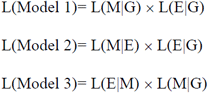

with G being the genetic variant, M the CpG site, E the gene expression, and L(Y|X) the likelihood of the standard linear model, with Y as the dependent variable and X as the predictor.

We then calculated each model’s probability using a uniform distribution of the priors.

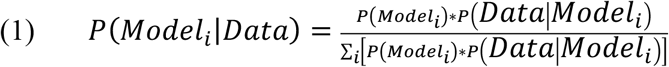

where *P* represents the probability of model i, and P(Model_1_)= P(Model_2_)= P(Model_3_)=1/3. The equation (1) can then easily be simplified as:

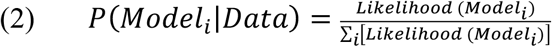

We calculated the probability of each model for all trios, and assigned each trio to the model presenting the highest probability, which we required to be higher than 0.9. If no models reached such a probability, the trio was declared insignificant. We found that reverse causation was indeed highly unlikely: at the non-stimulated state, only 3.1% of the trios were assigned to *Model 2*, while <1% of the trios were assigned to *Model 2* in the presence of immune stimulation.

### Supplementary Note 2

We found that the extent of sharing of eQTMs between individuals of African and European ancestry was significantly higher than that of meQTLs. To check that this observation is not explained by differences in power between the two analyses, we declared as “shared” all gene-CpG pairs (for the eQTM mapping) and all CpG-SNP pairs (for the meQTL mapping) that were detected in one population (FDR=5%) and whose *P*-value of association was lower than 0.05 in the other population. Among the 1,108 gene-CpG pairs detected in at least one population at FDR=5%, we found 708 pairs (63.4%) that were shared between AFB and EUB. In the meQTL mapping, among the total of 2,553,078 CpG-SNP pairs detected in at least one population at FDR=5%, we detected 1,003,271 pairs (39.3%) that were shared across populations. The level of population sharing was thus significantly higher for eQTMs than for meQTLs (OR ~2.5, Fisher’s exact *P* = 3.5×10^−51^), indicating that power disparities between the two analyses cannot explain the overall differences observed.

